# A neural network model for the evolution of reconstructive social learning

**DOI:** 10.1101/2024.09.18.613615

**Authors:** Jacob Chisausky, Inès Marguerite Daras, Franz J. Weissing, Magdalena Kozielska

## Abstract

Learning from others is an important adaptation. However, the evolution of social learning and its role in the spread of socially transmitted information are not well understood. Few models of social learning account for the fact that socially transmitted information must be reconstructed by the learner, based on the learner’s previous knowledge and cognition. To represent the reconstructive nature of social learning, we present a modelling framework that incorporates the evolution of a neural network and a simple yet biologically realistic learning mechanism. The framework encompasses various forms of individual and social learning and allows the investigation of their interplay. Individual-based simulations reveal that an effective neural network structure rapidly evolves, leading to adaptive inborn behaviour in static environments, pure individual learning in highly variable environments, and a combination of individual and social learning in environments of intermediate stability. However, the evolutionary outcome depends strongly on the type of social learning (social guidance versus social instruction) and the order of individual and social learning. Moreover, the evolutionary dynamics of social learning can be surprisingly complex, with replicate simulations converging to alternative outcomes. We discuss the relevance of our modelling framework for cultural evolution and suggest future avenues of research.

## Introduction

Learning is an adaptation that allows individuals to modify their behavioural phenotype to better fit their environment^1–3^. As such, learning requires an input of information relevant to the current state of the environment. This information may come from one of two primary sources: directly from the environment or indirectly from other individuals. If an individual relies exclusively on their own interaction with the environment without any social influence, we talk about individual learning. For instance, an animal may sample various food items from the environment to assess their quality and modify its future foraging based on the gained knowledge (trial-and-error learning). Information derived from other individuals is utilised in social learning, in which the behaviour of another individual affects the learning process. This social influence could take many forms, from simply making the learners attend to specific food items when observing others foraging (so-called stimulus enhancement) to the active process of information transfer from one individual to another (teaching)^4,5^.

Social learning is the cornerstone of cultural evolution as it allows the transfer of information and skills between individuals^6,7^. While experimental and observational studies shed some light on the evolution of culture, theoretical models are important to understand and predict the longer-term implications of cultural transmission via social learning. An adequate representation of social learning in these models is crucial for a sound understanding of cultural evolution. Based on the idea that the spread of cultural information resembles the spread of genetic variants in genetic evolution or the spread of infectious agents in an epidemic, most cultural-evolution models resemble models from population genetics and epidemiology^6,8,9^. Many of these models assume, often implicitly, that ‘cultural traits’ are directly transmitted between individuals. This assumption has been criticised (e.g.^10–12^) because social learning fundamentally differs from the direct transfer of ‘cultural information’ from one individual to the other. As far as we know, learned information is not replicated and stored at a specific location in the brain but instead ‘reconstructed’ by the learner, based on the learner’s skills and knowledge and filtered by cognitive processes^13–15^. It has been argued that the mechanisms underlying social learning are only of marginal importance for the course and outcome of cultural evolution^13^. This claim can, however, only be judged when more realistic representations of social learning are incorporated into cultural evolution models. Here, we present a conceptual model of such a representation.

Our model of social learning is based on neural networks (see e.g.^16–19^ and Discussion for earlier studies using similar approaches). These networks allow the seamless integration of incoming information (from other individuals or the environment) with previous ‘knowledge’ and thus reflect the learner-specific ‘reconstruction’ of the learning content. Our model is based on previous work on the evolution of neural networks underlying individual learning^20^. In this baseline model, important aspects of the neural network are genetically heritable (and hence evolve by genetic evolution), while other, non-heritable aspects of the network change in the lifetime of an individual as a result of personal experience. The learning mechanism is biologically inspired (it resembles dopamine prediction error coding in the brain^21,22^) and affects only a small part of the neural network. To a large extent, the network is determined by genes, which, together with the genes encoding other aspects of the learning strategy (e.g. the timing and duration of learning), evolve and, consequently, adapt the system to the environment, leading to more efficient learning^20^.

Here, we expand this model for individual learning by including two forms of social learning, which we call ‘social guidance’ and ‘social instruction.’ Socially guided learning resembles social enhancement in which learners observe others (demonstrators) interacting with specific items from their environment and then learn by direct interaction with the same items. Socially instructed learning resembles teaching as learners do not interact with the environment directly but obtain information about the environment from the teacher^5^.

The proposed modelling framework has a number of desirable properties. First, learning is a reconstructive process^10–12,23^. ‘Knowledge’ is not stored at a specific location but distributed over the whole neural network. The learning process depends on both, the information offered and the state of the network. Therefore, different individuals will draw different conclusions from the same information, due to differences in their inherited network and/or differences in their learning history. Second, individual and social learning utilise a common underlying mechanism^14,24^.

Depending on the type of information received, the neural network adapts via individual learning, social learning, or a combination of both types of learning. Therefore, one basic model allows us to study (the evolution of) different forms of individual and social learning and any combination thereof. Third, the modelling framework integrates genetic adaptation of the neural network and individual adaptation to the prevailing circumstances via learning. The integration of both processes yields new perspectives on gene-culture coevolution^25^ (which are beyond the scope of the current study). Fourth, the proposed modelling approach can be applied to a broad variety of learning tasks. Here, we present the modelling framework and study some of its basic properties. We keep the neural network small and simple in order to demonstrate that efficient (social) learning can emerge even in a simple configuration. We keep the network architecture fixed, although it could easily be made evolvable. Perhaps most importantly, we focus on a simple learning task: learning the quality of (food) items on the basis of external cues (like colour or smell), where the relationship between cues and quality may change over the course of time. Using the same learning task and network structure as in our previous study on individual learning^20^ allows us to compare the impact of individual learning, social learning, and their interplay.

Throughout, we systematically change the properties of the environment, most importantly the rate and degree of environmental change over time. Based on intuition and earlier work on social learning^7,26,27^, we expect that neither individual nor social learning will evolve in stationary environments, that pure individual learning will evolve in highly variable environments (where the information acquired by the previous generation gets soon outdated), and that social learning (or a mixture of social and individual learning) will evolve in environments with intermediate stability. By means of individual-based simulations, we will show that, by and large, the model outcome is in line with these predictions. However, the outcome depends strongly on the type of social learning (social guidance versus social instruction). Moreover, the evolutionary dynamics of social learning can be surprisingly complex, with replicate simulations converging to alternative outcomes.

## Model overview

We consider a population of individuals that live in an environment with characteristics that remain stable throughout an individual’s lifetime but change over longer time periods. The individuals have the task of collecting food of high quality. Food quality cannot be perceived directly, but it is associated with cues like colour or smell. Each individual harbours a simple neural network that allows it to predict the quality of a food item on the basis of its associated cue. The neural networks are genetically encoded and transmitted from parents to their offspring, subject to some rare mutations. Hence, the networks can evolve over the generations, providing newborn individuals with an ‘inborn’ knowledge of the association between cues and quality. As the environment can change (Supplementary Fig. 1), this knowledge can become outdated. To cope with this, individuals can evolve the ability to spend some time learning at the beginning of their lifetime. During this learning period, an individual can spend a number of learning episodes to sequentially test food items by comparing the predicted quality *P* of each item with a ‘target value’ *T* (see below). If there is a discrepancy between *P* and *T*, the neural network of the individual is changed by making use of the ‘Delta rule,’ a simple and biologically realistic learning algorithm that acts at the level of individual neurons^20^. These changes in the network due to learning are not transmitted to the offspring.

Figure 1 illustrates the four types of learning considered in this study. Each learning event and its outcome depend on two aspects, the choice of items tested during learning and the target value on which the feedback to the learning individual is based. Both aspects can depend on individual and social information. Kozielska and Weissing^20^ considered an implementation of individual learning where the food items to be learned from are chosen at random and learning is based on the discrepancy between the food item’s true quality and the quality of the item predicted by the learner (left panels in Fig. 1). We call this form of learning <u>‘unguided individual learning’</u>. For the purpose of this study, we will focus on <u>‘self-guided individual learning’</u> (second column of panels in Fig. 1), which differs from unguided learning in one aspect: the item chosen for learning is the ‘most promising’ from a random set of items, that is, the item with the highest predicted quality. We also consider two variants of social learning, where learning is influenced by a ‘demonstrator’ (a randomly selected individual from the parental generation). <u>‘Socially guided learning’</u> differs in one aspect from self-guided learning: the item chosen for learning is pointed out by the demonstrator, who recommends the item for learning that, according to the demonstrator’s neural network, has the highest predicted quality. Finally, we consider <u>‘socially instructed learning’</u> (right-most column of panels in Fig. 1), which differs from socially guided learning in that learning is not based on the discrepancy between the true quality and the quality predicted by the learner, but on the discrepancy between the demonstrator’s and the learner’s prediction of quality. Hence, in the first three variants of learning, the target value is the true quality of the item to be learned from, while in the case of socially instructed learning, the target value is the demonstrator’s prediction of the item’s quality. Accordingly, socially instructed learning misses the ‘reality check’ that is part of the other forms of learning, and the learning outcome strongly depends on how knowledgeable the demonstrator is about the true state of the environment. On the positive side, the lack of a reality check may speed up socially instructed learning. Therefore, we will later consider the possibility that socially instructed learning occurs faster than the other types of learning. Additionally, combining individual and social learning within an individual’s learning period adds a reality check to socially instructed learning. It may be advantageous to combine individual and social learning in a ‘strategic’ way. Therefore, we will later allow for the evolution of a ‘learning schedule’ where social learning can be utilised before individual learning or the other way around. The value of social learning can also be enhanced by learning from the most knowledgeable individuals^28,29^. Therefore, we later implement a ‘success bias’ in demonstrator selection, so that members of the parental generation who achieved a higher lifetime energy gain (a proxy for fitness) in their life are more likely to be selected as demonstrators.

**Fig. 1.**
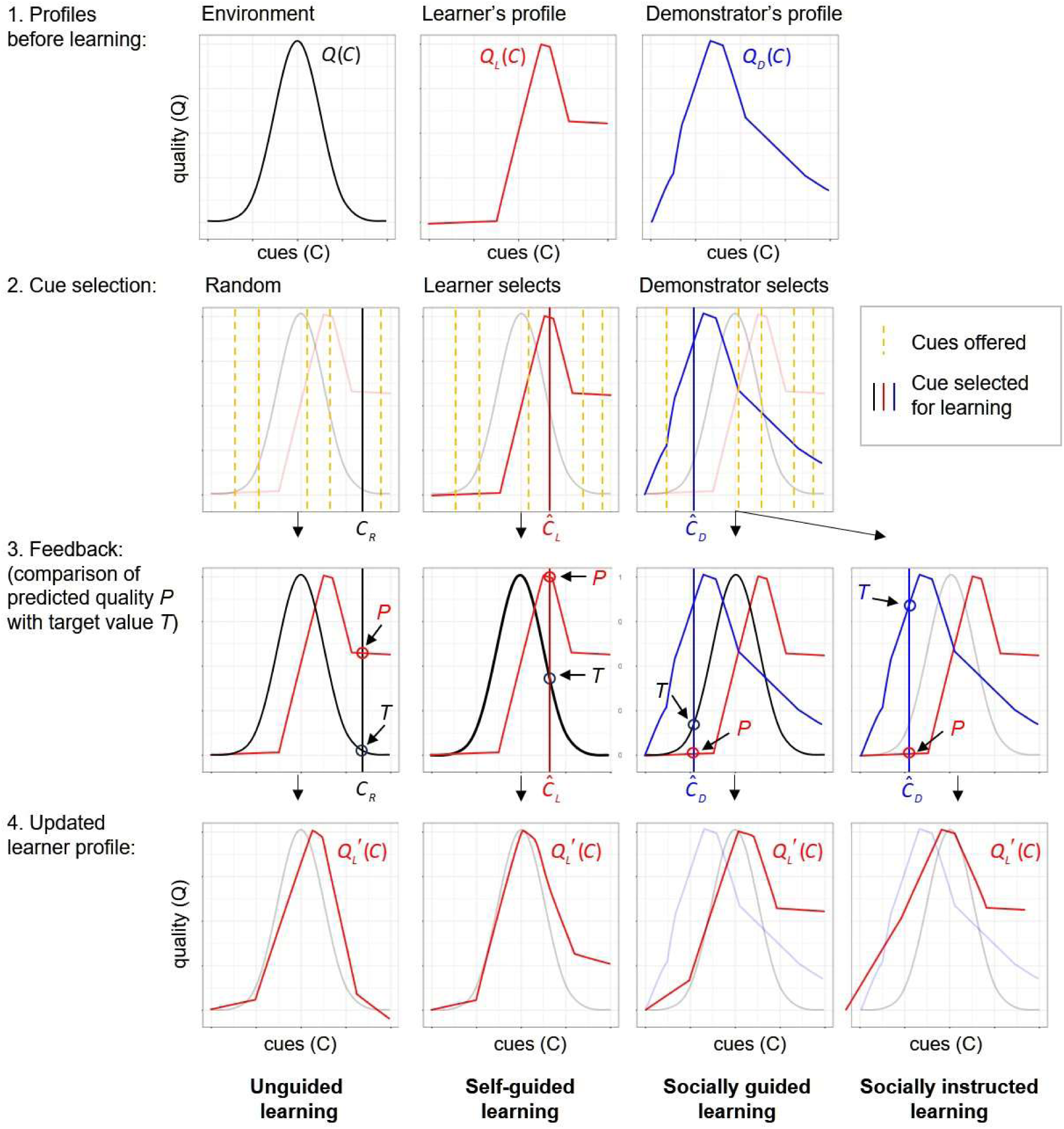
The four types of learning considered in our model. Individuals have the task of selecting the highest-quality food item among a set of available items. Food quality cannot be perceived directly, but only on the basis of cues like colour or smell. The ‘true’ relationship between cues and quality is described by the ‘environmental profile’, a function *Q*(*C*) that can change between generations. Each individual harbours a neural network that predicts the relationship between cues and quality. This network is inherited but can change during the individual’s lifetime due to individual or social experience, as described below. The coloured curves illustrate the profiles (= the predictions of quality from cues) of two individuals, a ‘learner’ (*Q_L_* (*C*), red curve) and a ‘demonstrator’ (*Q_D_* (*C*), blue curve). A learning event is modelled as follows: first, a number of random cues are offered to the learner (the six vertical lines in the second row of panels, though 10 are utilised in our simulations). Second, the learner selects a cue in one of three ways: at random (*C_R_*, left panel), the cue the learner predicts to be associated with the highest quality (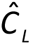, middle panel), or the cue the demonstrator predicts to be associated with the highest quality (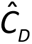 right panel). In all three cases, the learner has a prediction *P* of the quality associated with the selected cue 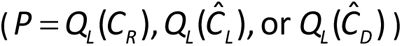. Third, the learner receives feedback on the correctness of the prediction by comparing it to the ‘target value’ *T*. In the first three types of learning, the learner is informed about the ‘true’ quality of the selected item 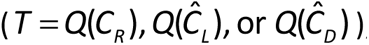; in the fourth variant, the learner is informed about the quality predicted by the demonstrator 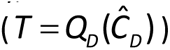. Fourth, the learner’s neural network is updated by applying the Delta rule (see Methods) that incorporates the discrepancy between predicted value *P* and target value *T*. This update leads to a changed prediction profile of the learner 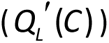, who uses this network as the starting point of the next learning episode or, after learning is complete, uses it to make foraging decisions.

Our model includes the basic ingredients of ‘<u>gene-culture coevolution’</u>. As indicated above, the model includes the social (‘cultural’) transmission of information. The setting in which this transmission occurs is strongly affected by heritable components and, hence, subject to evolution by natural selection. Not only the neural networks are heritable but also the parameters determining the learning process (such as the duration of the learning period and the learning rate in the Delta rule). We also allow the evolution of combined individual and social learning. In this case, the relative frequency of each type of learning and/or the scheduling of the two types of learning are also heritable properties.

Once the learning period is over, individuals switch to foraging where they use their neural network to assess the available foraging options. In each time step spent on foraging, they are offered a number of food items of which they choose the one which, based on the items’ cues, has the highest predicted quality. As the predictions of the network may not be perfect, one of the other items may actually be of higher quality. The ratio of the quality of the chosen item and the quality of the highest-quality item on offer is a measure of <u>network performance</u>. To be more precise, we quantify ‘performance’ by dividing the sum of the qualities of the items chosen throughout the foraging period by the sum of the maximal qualities on offer in this period. Therefore, performance equals one if, during the foraging period, an individual always chooses optimally, and it is smaller than one otherwise.

The sum of the qualities chosen throughout the foraging period (lifetime energy gain) determines the expected reproductive success (‘fitness’) of an individual. Those individuals with a higher fitness transmit their heritable properties to more offspring, eventually leading to the spread of neural networks and learning parameters that are best adapted to the characteristics of the environment. It is important to notice that there is a <u>trade-off between exploration and exploitation</u>^20^: the longer the learning (‘exploration’) period, the shorter the foraging (‘exploitation’) period, but typically the higher the average performance in each time step of the foraging period.

## Results

### Evolution of unguided and self-guided individual learning

The evolution of unguided individual learning was studied in detail in Kozielska and Weissing^20^. Here, we start by comparing the outcome of evolution for the two types of individual learning: unguided and self-guided learning. To this end, we switched off social learning by setting the proportion of social learning (*π_s_*) to zero.

Figure 2 shows the evolved number of learning episodes and the achieved performance for the two individual learning implementations (see Supplementary Fig. 4 for the corresponding lifetime energy gain, our proxy for fitness). By and large, the number of learning episodes is very similar for both implementations. In particular, learning tends to evolve under the same conditions: when the environment changes more frequently and/or the changes are larger. Although the simulations exhibit a clear pattern, when learning evolves, the number of learning episodes may differ considerably (often between 15 and 25 or even more) across replicate simulations. At least three factors can cause this variation. First, the inherent stochasticity of individual-based simulations makes each replicate unique. Second, replicates may differ in the evolved neural networks which may affect the number of learning episodes required for efficient learning. Third, the exact number of learning episodes seems to be under weak selection. Accordingly, this number can fluctuate considerably over evolutionary time within each simulation.

**Fig. 2.**
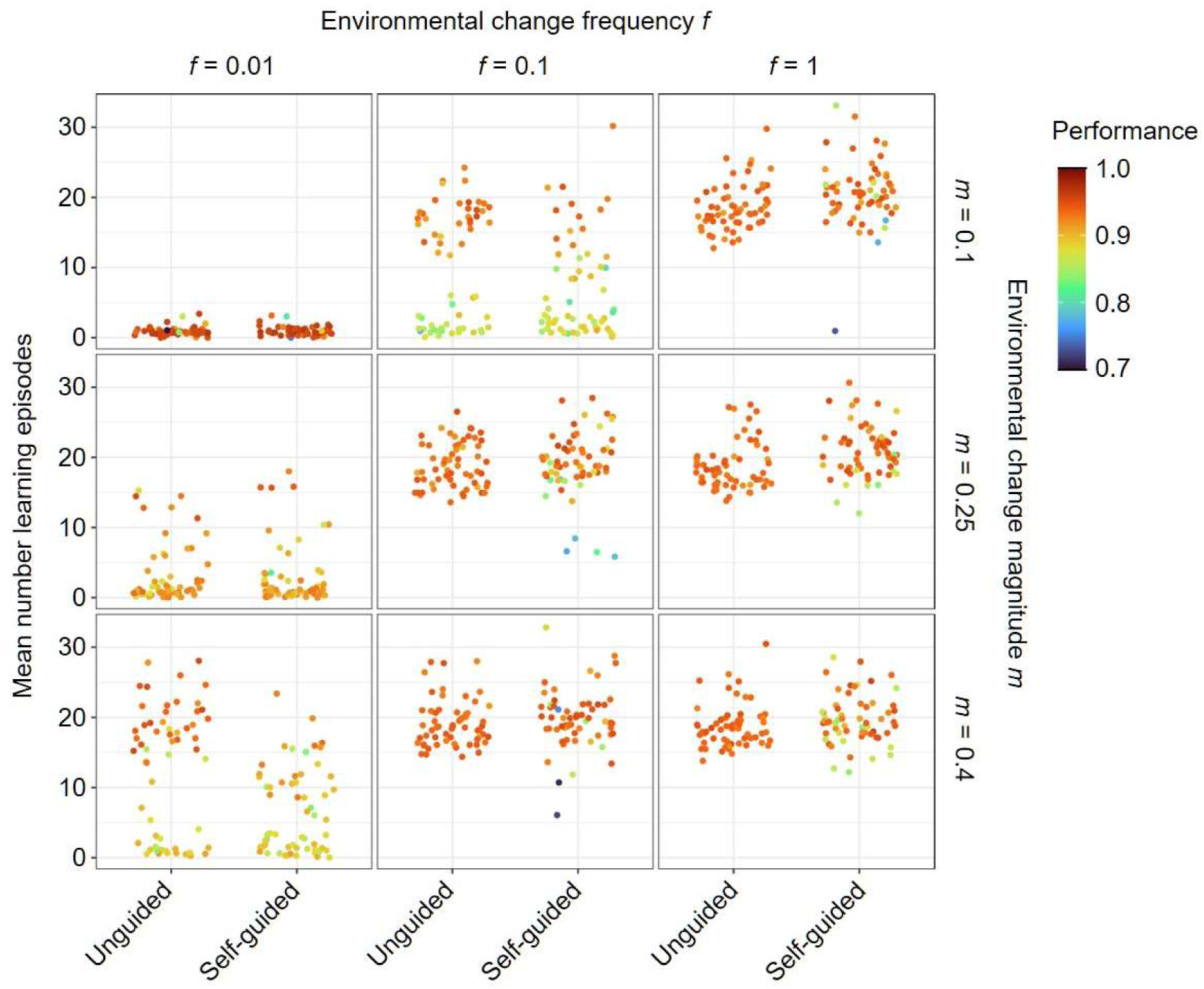
Evolved time spent on unguided or self-guided individual learning in relation to the frequency and magnitude of environmental change. For three frequencies (*f*) and magnitudes (*m)* of environmental change, the panels show the number of learning episodes that evolved within 20K generations in the cases of unguided and self-guided individual learning. A small number of learning episodes indicates that learning did not evolve. Each panel shows the outcome of 60 replicate simulations per learning implementation (unguided vs. self-guided individual learning). Each point represents the population mean averaged over the last 2K generations of one simulation. The colour of each point represents the average performance in the population (‘performance’ = the quality of food items chosen, relative to the maximal possible quality). Artificial noise is added to the *x*-axis to improve readability. In all simulations, the width of the environmental profile was σ = 0.25 and social learning was not allowed to evolve.

In both implementations of individual learning, populations achieve quite high performance. However, the performance achieved by self-guided learning is more variable between replicate simulations and on average it is somewhat lower than that achieved by unguided learning (Fig. 2). The width of the environmental profile (σ; see Methods) only slightly affects the evolution of learning, and the outcome of evolution for the two individual learning implementations is similar in most environments (but see Supplementary Figs 3 and 4).

In conclusion, different implementations of individual learning do not have a large impact on the evolved strategy (learning vs no learning). For the rest of the paper, we decided to focus on self-guided learning as it is more similar to our implementation of social learning in the sense that organisms do not randomly explore all possible cues but are guided by the preferences of the demonstrators. This way, we can make a more direct comparison between individual and social learning, without the confounding effect of random exploration of the environment when learning individually. However, the Supplementary figures show also the results for the coevolution of social learning with unguided individual learning and the general conclusions are usually the same as for self-guided learning.

### Evolutionary trajectories and classification of evolutionary outcomes

For the rest of the paper, we will use the self-guided implementation of individual learning, additionally allowing social learning to evolve. That means that in addition to the evolution of the total number of learning episodes (and individual learning rate), the proportion of social learning (and social learning rate) could also evolve. For most of the paper, we focus on analysing the evolved strategies by looking at the number of learning episodes and the proportion of social learning.

To illustrate the visualisation and interpretation of our simulation outcomes, we start with an example that will later be discussed in more detail (see Fig. 8). Figure 3 shows the evolutionary trajectories of four replicate simulations (identical parameter settings and initial conditions) demonstrating the variation in the course and outcome of the evolutionary process. In some simulations (e.g., Fig. 3A), the number of learning episodes converges to a very low value, meaning that the population has lost learning altogether. Once this has happened, the proportion of social learning does not matter anymore. Accordingly, this proportion is evolving via genetic drift, corresponding to a random walk along the vertical axis. In other populations (e.g. Fig. 3B), pure individual learning evolves: the number of learning episodes evolves to a value that is clearly higher than zero while the proportion of social learning evolves to a value close to zero. Another potential outcome is pure social learning with a positive number of learning episodes and a proportion of social learning close to one. However, this is usually a transient state (Fig. 3A, generations near 2K). Lastly, a combined individual and social learning strategy (in the literature sometimes called a mixed strategy) can evolve with an intermediate proportion of social learning (Fig. 3C). In some simulations, populations switch between different strategies (Fig. 3D). The legend to the right of Fig. 3 classifies the evolved learning strategies into four categories (note that the borders between the regions are somewhat arbitrary). Throughout the rest of the paper, we will use graphs with the same axes, though instead of looking at the whole evolutionary trajectory we will focus on the population averages at the end of the simulations, representing each replicate simulation by a single point. It should be noted that for the regions at the border of the figure, populations are more homogenous, while in the combined-strategy region, a given proportion of social learning (and the number of learning episodes) at the population level can be achieved in multiple ways: all individuals in the populations can be similar and show a learning strategy given by the population average, or individuals can differ between each other with some showing more (social) learning than others. A survey of a subset of our simulations showed that the individuals of the evolved populations did generally not differ in the number of learning episodes while the proportion of social learning used can be quite variable across individuals. However, we never saw a clear separation into distinctive strategies. Therefore, we feel confident that focusing on population means is justified.

**Fig. 3:**
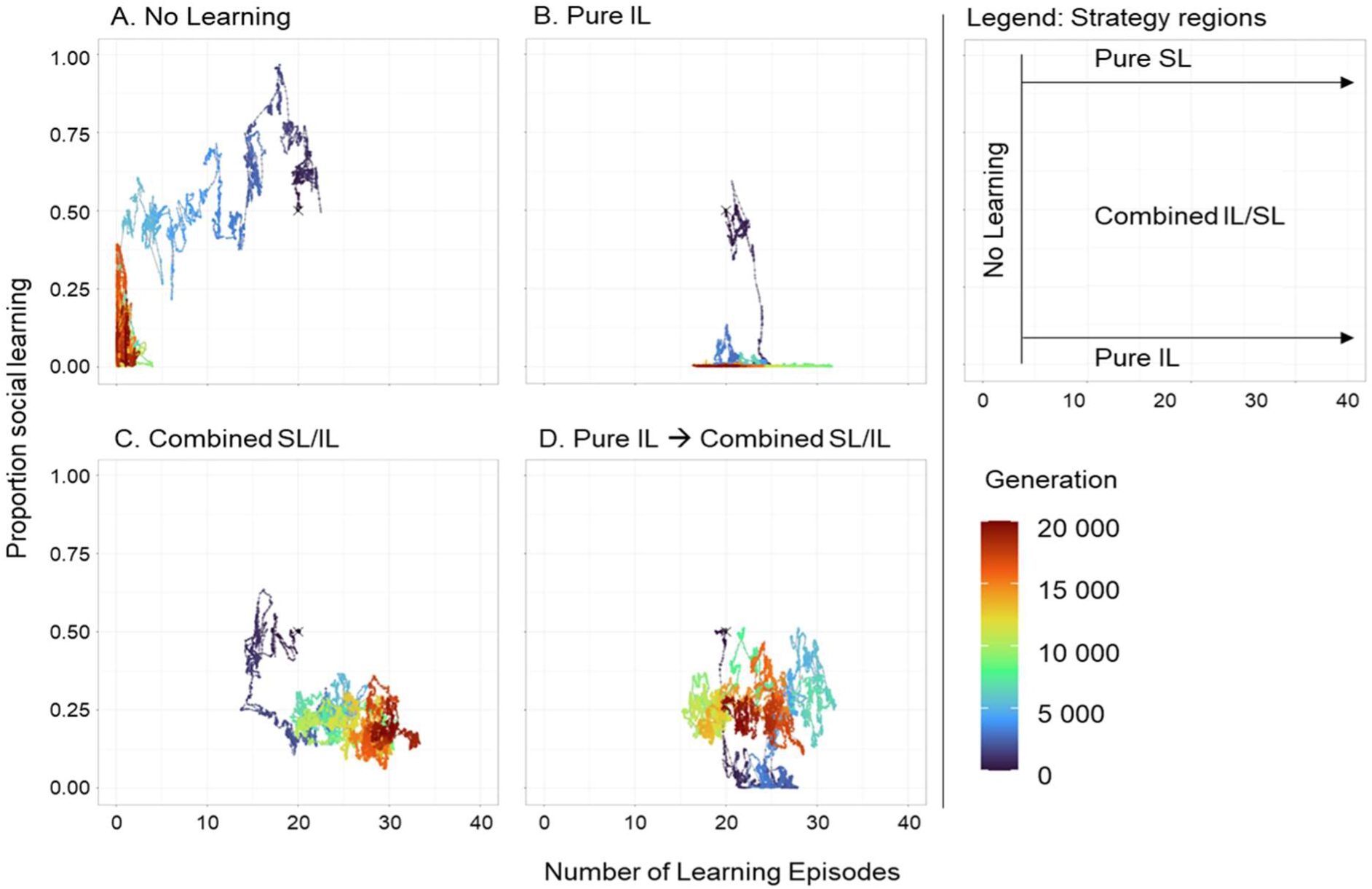
Evolutionary trajectories and alternative outcomes of evolution. Four replicate simulations illustrate that the co-evolution of individual and social learning can lead to markedly different outcomes. Evolutionary trajectories are coloured as indicated in the colour spectrum to the right: from dark blue in the initial generations to dark red towards the end of the simulation. The four trajectories all start with 20 learning episodes and a proportion of social learning of 0.5 (marked by a dark blue cross). After 20K generations (dark red part of the trajectories), learning was lost in replicate **(A)**, pure individual learning had evolved in replicate **(B)**, and a learning strategy combining about 20% of social learning with 80% of individual learning had evolved in replicate **(C)**. Replicate **(D)** illustrates that social learning, after having been lost initially, can later be regained, in this example resulting in a combination of 25% social learning and 75% individual learning. The legend to the right provides guidelines on how to interpret the simulation outcomes: ‘no learning’ if the evolved number of learning episodes is very low; ‘pure individual learning (IL)’ if the evolved proportion of social learning is very low; ‘pure social learning (SL)’ if the proportion of social learning is close to 1.0; and ‘combined individual and social learning (IL/SL)’ if the proportion of both types of learning is intermediate. The simulations were run for the same scenario as in Fig. 8, which will be discussed in detail later.

### Evolution of socially guided learning

We first investigate in which environmental conditions socially guided learning evolves, in which the demonstrator chooses a cue and the learner updates its network based on the quality of that cue. Figure 4 shows that for the most stable environments (*f* = 0.01 and *m* = 0.1) learning was lost in all simulations. For the least stable environments (*f* = 1 and *m* = 0.4) pure individual learning evolved in most replicates. In intermediate environments, a combination of individual and social learning evolved in many replicates. In some of these environmental regimes (*f* = 0.1 and *m* = 0.25 or 0.4 and *f* = 1 and *m* = 0.1), the combined strategy seems to be a strong attractor with (almost) all replicates evolving a similar combined strategy, reaching high performance. In contrast, in more stable regimes (e.g. *f* = 0.01 and *m* = 0.25 or 0.4 and *f* = 0.1 and *m* = 0.1) there is a considerable variation between replicates in the number of learning episodes, the proportion of social learning and the achieved performance. In some replicates, pure social learning evolved, but it was often linked with relatively low performance.

**Fig. 4.**
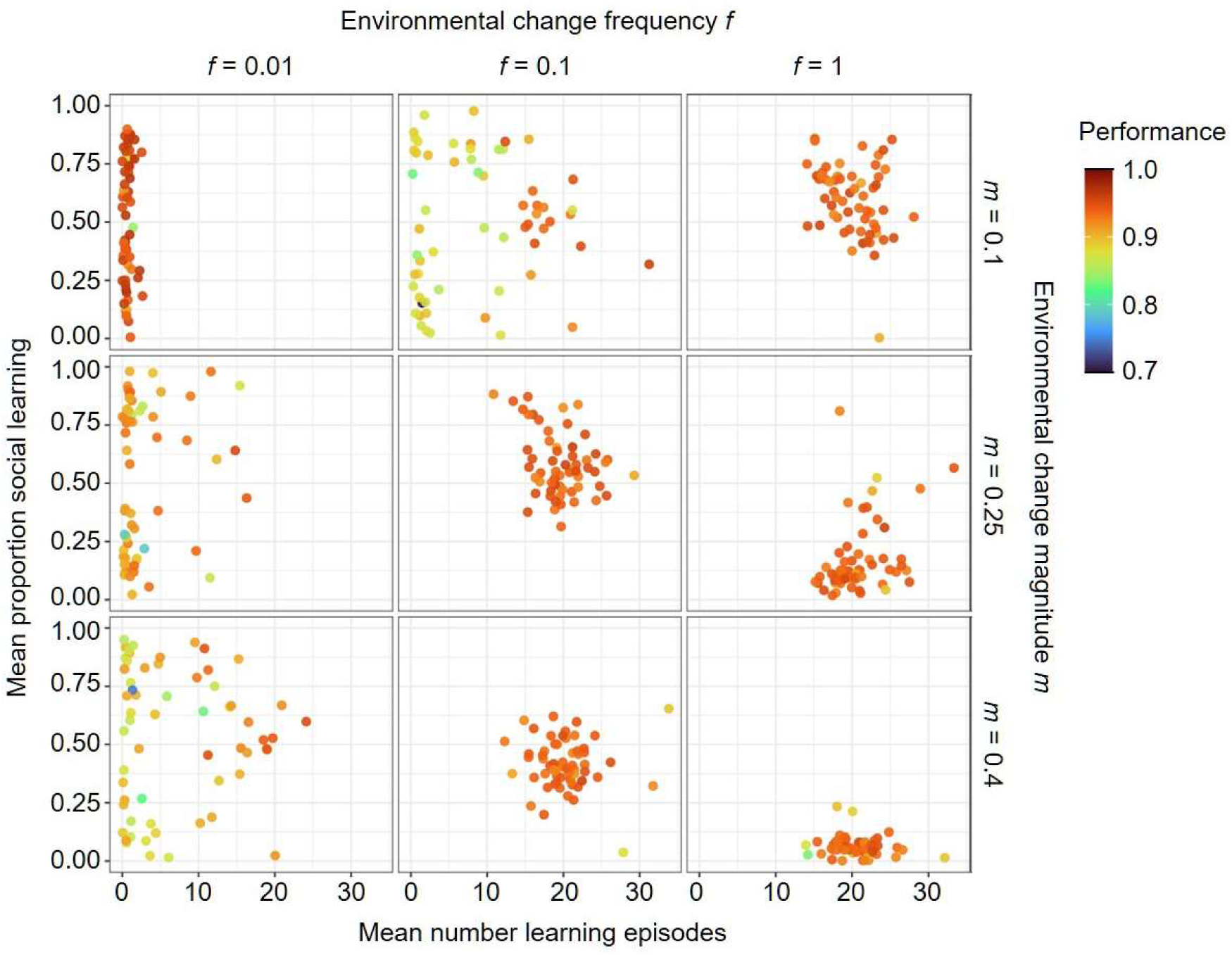
Joint evolution of self-guided individual learning and socially guided learning. For three frequencies (*f*) and magnitudes (*m*) of environmental change, the panels depict the characteristics of the evolved learning strategy after 20K generations when self-guided learning and socially guided learning could jointly evolve. Each panel shows the outcome of 60 replicate simulations. Each point represents the population mean number of learning episodes (*x*-axis) and mean proportion of social learning (*y*-axis), both averaged over the last 2K generations of the simulation. The colour of each point represents the average performance of the evolved learning strategy. A small number of learning episodes indicates that learning did not evolve. If learning evolved, a low proportion of social learning indicates the evolution of pure individual learning (here: self-guided individual learning), a high proportion indicates the evolution of pure social learning (here: socially guided learning), and an intermediate proportion indicates the evolution of a combined learning strategy. In all simulations, the width of the environmental profile was σ = 0.25.

Interestingly, when learning evolved, replicates with the highest performance were the ones where the number of learning episodes was around 20, independent of the environmental regime (similar to Kozielska and Weissing^20^). On the other hand, the proportion of social learning seems to be more affected by the environment. When learning evolved, the proportion of social learning decreased with increasing magnitude and frequency of environmental change. The results are only slightly affected by the width of the environmental profile, σ (see Supplementary Fig. 5).

In conclusion, socially guided learning evolves under a wide range of environmental conditions and predominantly as part of a strategy that combines individual and social learning. The results in Fig. 4 are generally in line with the expectations stated in the Introduction: less stable environments lead to less social learning and more individual learning, but in very stable environments learning does not evolve at all.

Socially guided and self-guided learning only differ regarding the cues individuals choose to learn from. A wide prevalence of a combination of socially guided and self-guided learning indicates that it is often beneficial to rely on two sources of information (other individuals’ and own preferences) with respect to which part of the environment to focus on when learning. This benefit does not only stem from the fact that this way potentially a wider range of environmental cues is explored, as the results of socially guided learning paired with unguided learning are similar to those for self-guided individual learning (see Supplementary Figs 5 and 6).

### Evolution of socially instructed learning

A similar analysis to the one described above was performed for the situation where socially instructed learning could evolve together with self-guided individual learning. In socially instructed learning the demonstrator not only chooses the cue to learn from but also provides the learner with the expected quality of that cue (i.e. the learner has not access to the real quality of the cue when learning socially). Figure 5 clearly shows that now social learning is selected against in all environmental regimes. No learning evolves in stable environments, while pure individual learning evolves in less stable environments. Selection against socially instructed learning is present for all considered widths of the environmental profile (*σ*) and also when it co-evolves with unguided individual learning (Supplementary Figs 7 and 8).

**Fig. 5.**
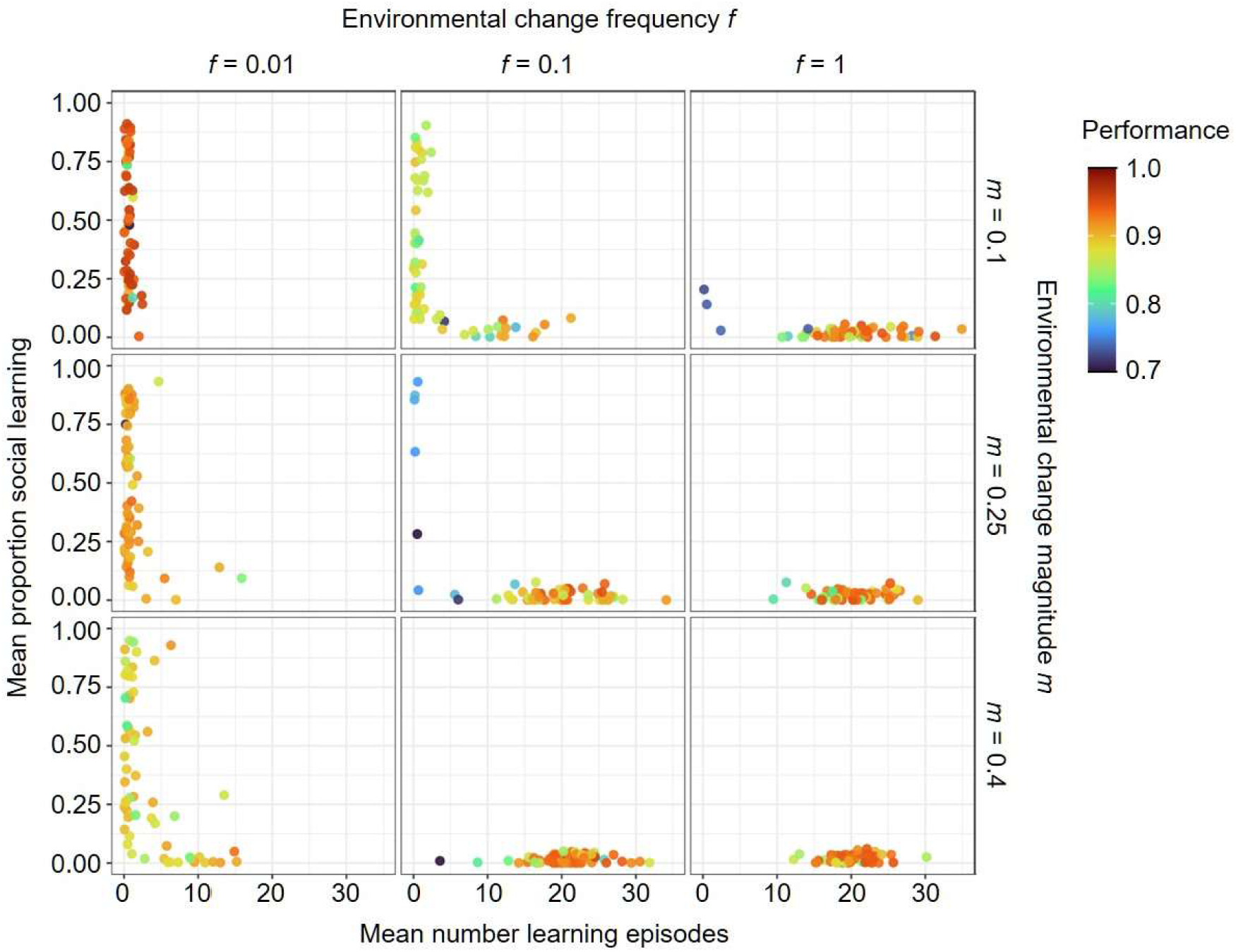
Joint evolution of self-guided individual learning and socially instructed learning. For three frequencies (*f*) and magnitudes (*m*) of environmental change, the panels depict the characteristics of the evolved learning strategy after 20K generations when self-guided learning and socially instructed learning could jointly evolve. Each panel shows the outcome of 60 replicate simulations. In all simulations, the width of the environmental profile was σ = 0.25. Plotting conventions as in Fig. 4.

Why is socially instructed learning selected against? A disadvantage shared with socially guided learning is that the ‘beliefs’ of the demonstrators regarding the location of the peak of the environmental profile may be outdated because the environment has changed between generations. However, there is a second disadvantage that may be more decisive. While in socially guided learning learners can compare their beliefs on the quality associated with the chosen cue with the true quality, such a ‘reality check’ is missing in the case of socially instructed learning, where feedback is not provided from the true quality but from the demonstrator’s belief regarding the quality associated with the chosen cue. The demonstrator’s beliefs may not be precise enough to be a good basis for learning, especially since learning by itself is not an error-free process.

### Success bias, speed of learning and learning schedule

So far in our model, individual and social learning differed only in the information they provided but we did not take into account some properties of social learning that are often considered beneficial or necessary for social learning to evolve. These include success bias, the higher speed of social learning and the potential advantages of scheduled learning. We tested how the results of our model for instructed learning are affected if our model is expanded by implementing the above.

To incorporate success bias, each time a learner uses social learning, it chooses a demonstrator not randomly but with a chance proportional to that individual’s lifetime energy gain (i.e., ‘fitness’) in the previous generation. We did not want to incorporate an arbitrary cost to individual learning, so instead we allowed social learning to be ‘faster’ than individual learning, a property dubbed ‘SL Speed’. That means that in each learning episode of social learning, learning occurs multiple times. That is, an individual can interact with several cues and adjust their network more than once in the same time as is needed for one individual learning event. Lastly, we allowed a learning schedule to evolve. Until now, individual and social learning were combined in a random order (‘shuffled learning’). In the new simulations, this was still an option, but in addition, an ‘SL→IL schedule’ or an ‘IL→SL schedule’ could arise by mutation. In the former, the social learning episodes precede the individual learning episodes, while in the latter individual learning precedes social learning. We ran a new set of simulations in which success bias was either present or not, the speed of social learning was set to a value of 1 (as in earlier simulations) or 10, and the learning schedule could evolve or not (see Methods for the implementation of these extensions). This way we investigated the effect of all these factors in isolation and in combination with each other (Fig. 6). For these simulations, we used the environmental regime with intermediate stability (*f* = 0.1 and *m* = 0.25).

**Fig. 6.**
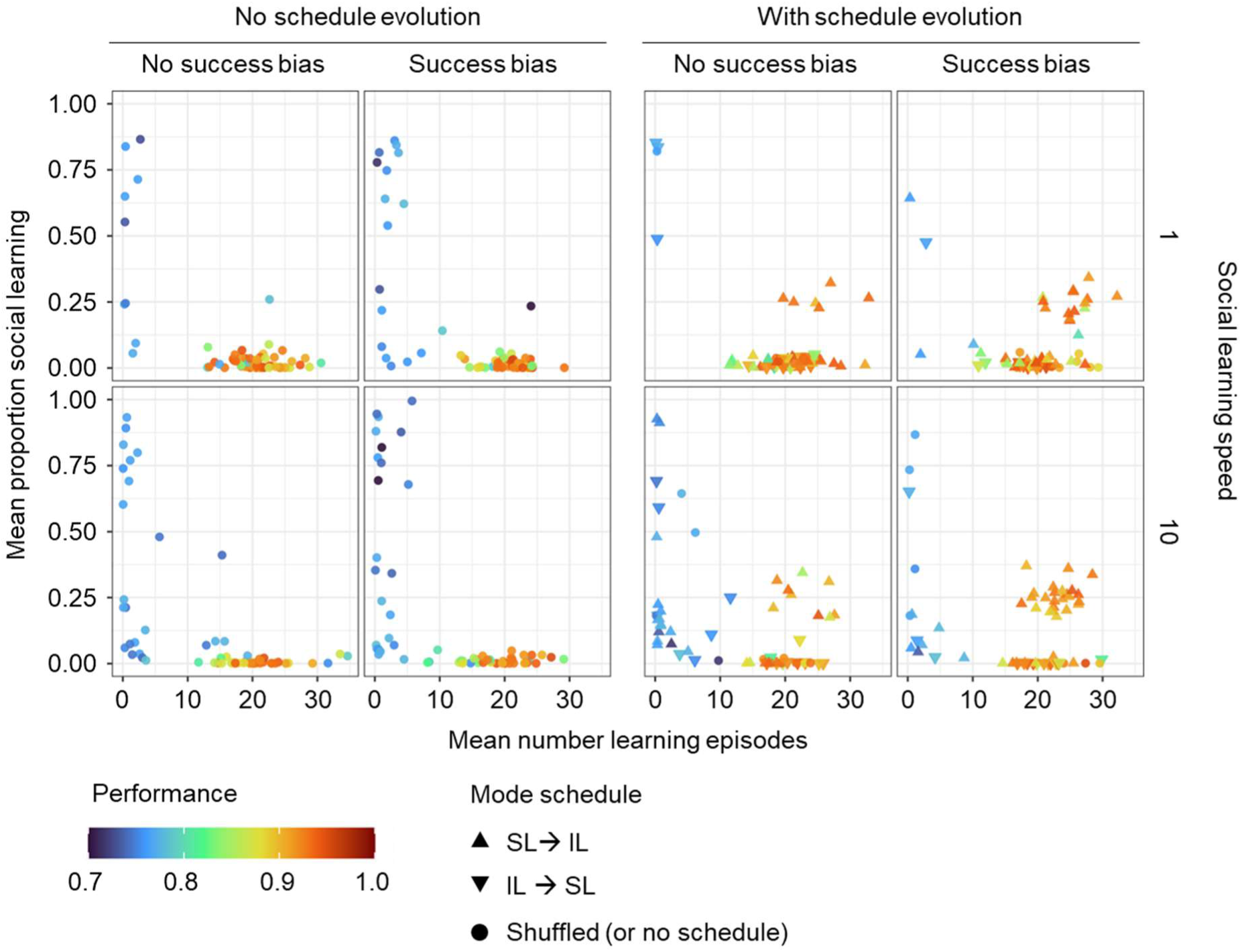
Effect of success bias, learning speed and learning schedule on the evolution of socially instructed learning. Evolved population mean number of learning episodes and the proportion of social learning (x- and y-axis, respectively) and the corresponding performance (colour) for 60 replicate simulations for each parameter setting (one data point corresponds to one replicate). Two left panels: All individuals used the ‘shuffled’ learning schedule, and the schedule could not evolve. Two right panels: The learning schedule could evolve. The most common schedule evolved in the population is indicated by the shape of the data point. Columns marked as ‘No success bias’ correspond to simulations in which learners chose demonstrators randomly. Columns ‘Success bias’ correspond to simulations in which learners chose demonstrators based on their lifetime energy gain as a proxy for success. Each row corresponds to different social learning speeds relative to individual learning (i.e. how much faster social learning is relative to individual learning). Environmental conditions are: frequency of change *f* = 0.1, magnitude of change *m* = 0.25 and width of the environmental profile σ = 0.25.

Figure 6 shows that adding success bias or increased SL speed by themselves or together is not enough to promote the evolution of socially instructed learning. The increased speed of SL seems to even lead to a lower probability of learning to evolve at all. This may be a result of our initial conditions in which networks are neither adapted to environmental challenges nor learning while at the same time the social learning proportion is relatively high (50%). In a situation like this, the mismatch between the demonstrators’ beliefs and the true state of the environment may lead to poor learning results, especially when the speed of social learning is high, and poor learning results may select for a loss of learning. Once lost, learning may be difficult to re-evolve as essential learning parameters (learning rates, proportion of social learning) are no longer under selection and can drift to values which inhibit learning.

While success bias and social learning speed by themselves were not favourable for the evolution of socially instructed learning, this form of social learning could get established as part of a combined strategy (with around 25% of the learning period spent on social learning) when the SL→IL schedule (social learning followed by individual learning) did evolve. Fig. 6 shows that such a combined strategy could evolve in the absence of a success bias or a higher social learning speed. However, high SL speed combined with success bias leads to the evolution of socially instructed learning in considerably more replicates compared to the situation without them, and the two factors seem to have a synergistic effect. This suggests that when individuals can learn from good demonstrators, the additional learning opportunities due to the higher SL speed are utilised well, making the SL→IL schedule even more beneficial. However, as seen in the simulations without schedule evolving, higher SL speed leads to a higher number of replicates without learning at all. In conclusion, a learning schedule with self-guided individual learning following socially instructed learning (SL→IL) is necessary for instructed learning to evolve and increased SL speed together with success bias may further facilitate this process.

Next, we looked at the evolution of instructed learning in different environmental regimes for the case of an evolving learning schedule, with success bias present and social learning speed set to 10.

Similarly to socially guided learning, socially instructed learning (given success bias, higher SL speed and evolution of learning schedule), is more likely to evolve in environments of intermediate stability (compare Fig. 7 with Fig. 4). However, in general, socially instructed learning is less likely to evolve than socially guided learning even with SL→IL schedule, success bias and higher SL speed. Instructed learning evolves in fewer environmental regimes and fewer replicates within a specific regime.

**Fig. 7.**
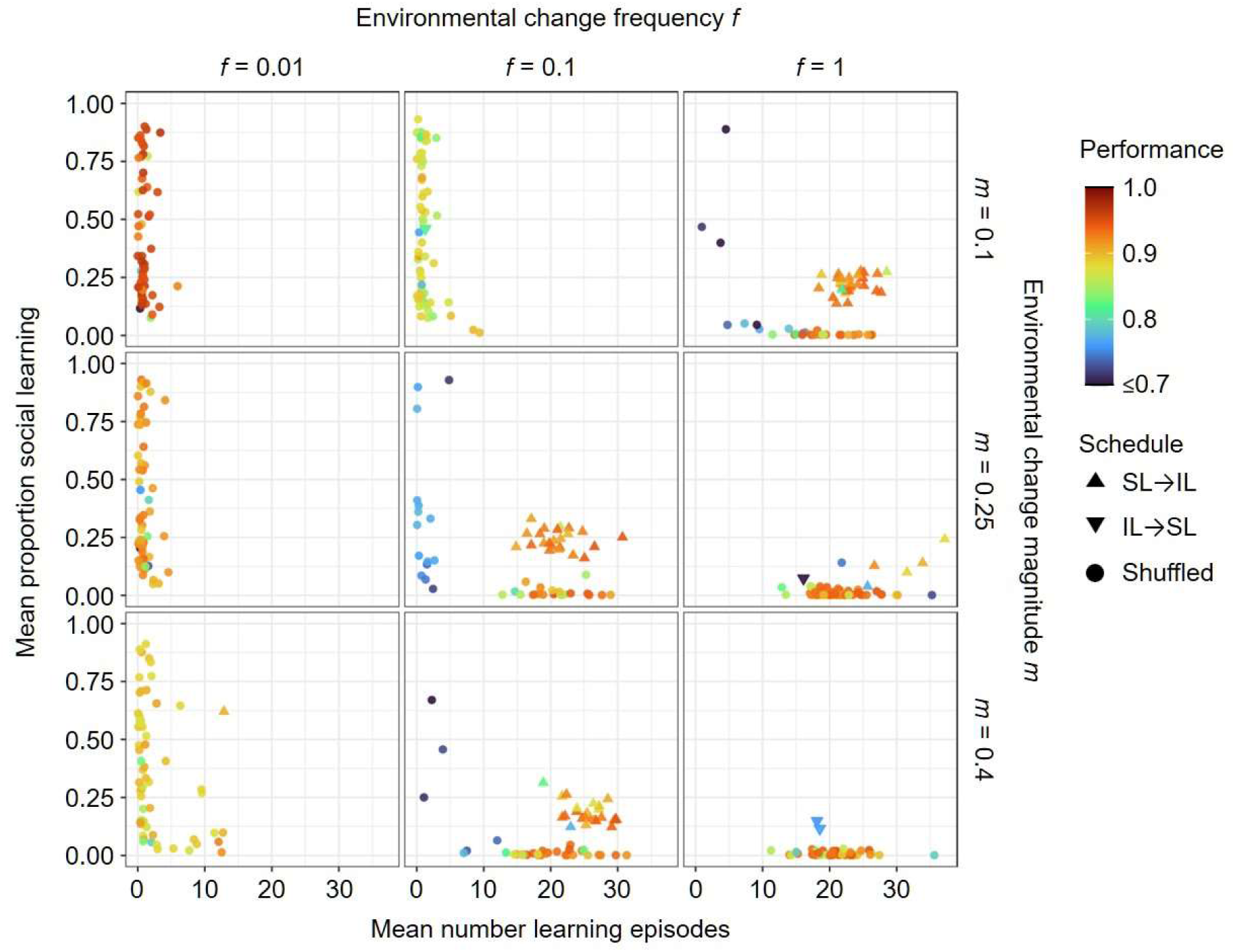
Joint evolution of self-guided individual learning and socially instructed learning with success bias, higher social learning speed and an evolving learning schedule. In these simulations SL speed was set to 10 and success bias was present. All simulations were initialised as ‘shuffled learning’, but the two learning schedules ‘SL→IL’ and ‘IL→SL’ could arise by mutation. Plotting conventions are as in Fig. 6.

As in Fig. 6, if socially instructed learning evolves, it is almost always associated with the SL→IL schedule (social learning proceeds individual learning). In the few exceptional cases where social learning evolved with a different schedule, the fitness achieved by these populations is clearly lower, indicating that these forms of learning correspond to a suboptimal strategy. Socially instructed learning can evolve also for narrower and wider environmental profiles (Supplementary Fig. 9).

However, social learning is less likely to evolve for the wide environmental profile with *σ*=0.4. Socially instructed learning is even less likely to evolve when unguided individual learning is utilised (Supplementary Fig. 10).

All in all, instructed learning can evolve as part of a combined strategy with SL→IL schedule in environments of intermediate stability. However, it seems that this strategy is not easily obtained or the attractor is not very strong as not all of the populations in a given environmental regime reached this evolutionary outcome.

### Fitness landscape, alternative attractors, and evolutionary transitions

In environmental regimes of intermediate stability we saw a large variability in evolutionary outcomes between replicates. A closer look at the evolutionary trajectories also showed that occasionally the predominant learning strategy within a population changes over the course of evolution, switching between pure individual learning, a combination of individual and social learning, and no learning at all (see Fig. 3D for an example).

To better understand the dynamics of the system and the observed outcomes, we analysed the evolutionary trajectories of 1000 replicate populations evolving in an environment of intermediate stability (frequency of change *f* = 0.1, magnitude of change *m* = 0.25). All individuals in all the replicates started using the same learning strategy (20 learning episodes, proportion of social learning 0.5, shuffled schedule) but with different random network weights and learning rates.

Again, we considered the combination of self-guided individual learning and socially instructed learning. The SL speed was set to 10, success bias was present, and a learning schedule could evolve. As shown in Fig. 8, we considered a phase space consisting of the number of learning episodes on the x-axis and the proportion of social learning on the y-axis. This phase space was divided into partitions of width 0.01 on each axis. For each simulation and each 101^st^ generation (after an initial period of 2K generations), the population average of lifetime energy gain (our proxy for fitness) was calculated and ascribed to the partition of phase space that was ‘visited’ by the population in that generation. This way, we could calculate the average fitness associated with each partition. Blank areas in the plot indicate that no population ended a generation in that region of parameter space.

**Fig. 8.**
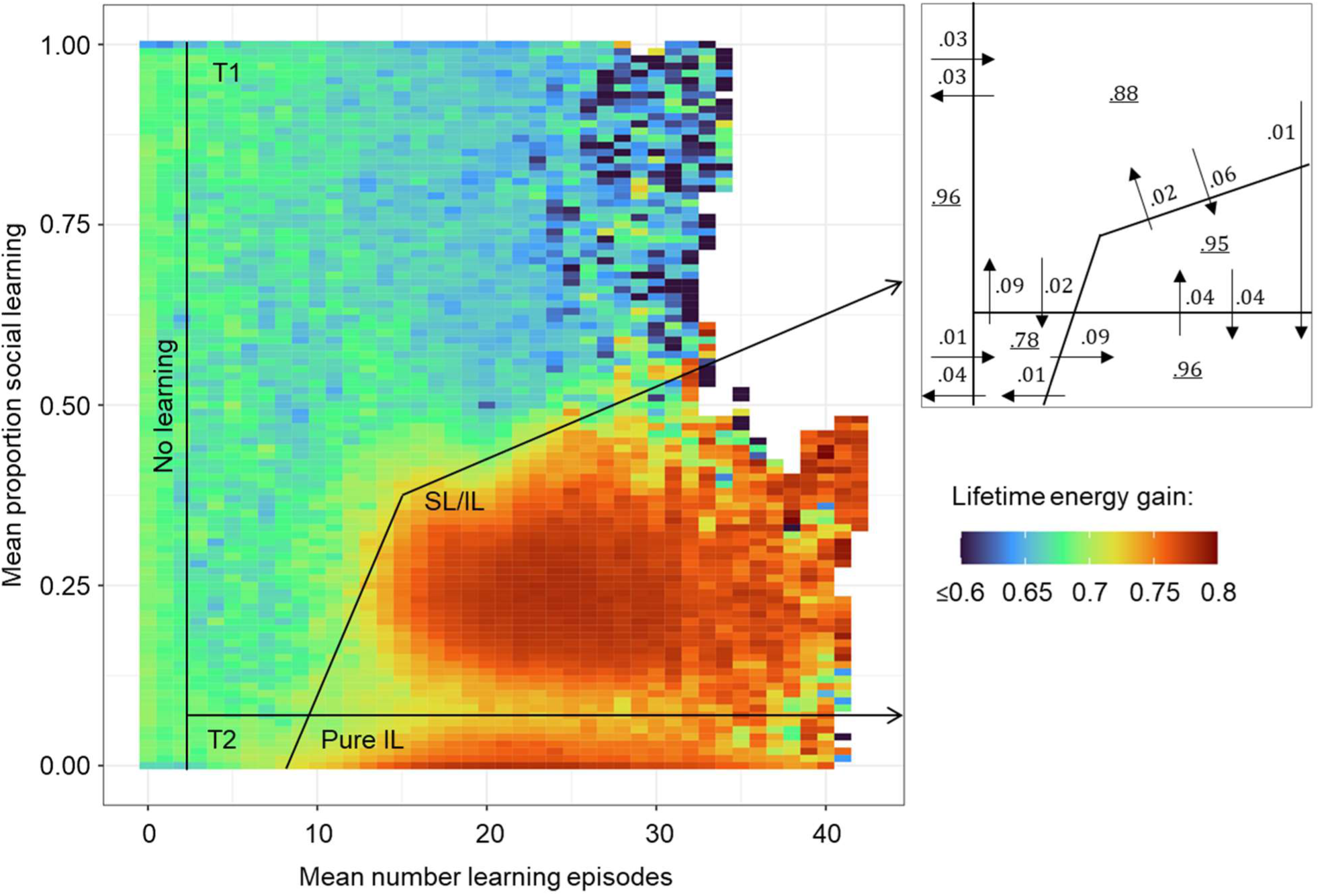
Fitness landscape for the joint evolution of self-guided individual learning and socially instructed learning. Heatmap representing average lifetime energy gain (a proxy for fitness; in the colour scheme expressed as a fraction of the theoretical maximum of 500) for evolutionary trajectories ‘visiting’ a given learning strategy (combination of the number of learning episodes and the proportion of social learning). Based on the heatmap and the evolutionary trajectories, we distinguish between five regions in phase space: no learning, pure individual learning (‘pure IL’), a high-fitness combination of individual and social learning (‘SL/IL’), and two ‘transitory’ regions T1 and T2. The scheme to the right indicates the probability and direction (arrows) of transition between strategy regions after 101 generations and the probability of being in the same region after 101 generations (number inside a region). The figure is based on the data from 1000 replicate populations evolving for 20K generations under intermediate environmental stability (*f* = 0.1, *m* = 0.25, σ = 0.25) with SL speed set to 10, success bias present and learning schedule evolving. The first 2K generations were not used as they represent the start-up phase when networks are not yet adapted to the environment and the learning task. (Note: The data for every 101^st^ generation, rather than every 100^th^ generation, was used to ensure that all generations before or after an environmental change were evenly sampled.)

Figure 8 shows a heatmap of all the data collected from these simulations. The ‘transition scheme’ to the right of Fig. 8 shows that there are three attractors: no learning resulting in low fitness and two high-fitness regions corresponding to pure individual learning (‘pure IL’) and a combination of individual and social learning (‘SL/IL’). Both types of learning have a similar fitness (though the combined strategy is marginally better; Supplementary Fig. 11). The two types of learning are separated by a shallow ‘fitness valley’. Both regions are relatively large showing that a broad set of parameter combinations leads to good outcomes. Populations transition between these regions at the same rate and when simulations are run for much longer times, both strategies are present among the replicates (Supplementary Fig. 12) but not necessarily in the same simulations.

The third evolutionary attractor from our simulations (‘no learning’) clearly has a lower fitness and is surrounded by two ‘transitory regions’: T1 with a considerable proportion of social learning and even lower fitness and T2 with a low proportion of social learning and similar fitness. Once a population loses learning, regaining it becomes relatively difficult. Once the number of learning episodes evolves towards 0, other learning parameters (learning rates, proportion of social learning and schedule) are no longer under selection and can drift, making it difficult for the population to evolve a better strategy, especially if the proportion of social learning drifts to become high or the IL→SL schedule is predominant in the population. The transitions between attractors highlight the fact that the learning strategy can change over time, even without a change in external evolutionary pressure.

### Evolved learning rates

While not the main focus of this paper, learning is also affected by the learning rates that are used in the Delta rule that governs the change in the network during a learning event. The value of the learning rate coevolves with the number of learning episodes and therefore is affected by environmental change ^20^. Here, we only briefly analyse the effect of the joint evolution of individual and social learning on the learning rates of the two types of learning. In the following simulations, we fixed the number of learning episodes to 25 and the proportion of social learning to 0.25 (a common outcome of evolution), as variation in these factors would confound the selection pressures on the learning rate. In addition, we analysed the effect of three fixed schedules: SL→IL, shuffled and IL→SL. To investigate the effect of the social learning speed we run the simulations with this parameter fixed to either 1 or 10. We run the simulations for socially instructed learning with success bias present.

In agreement with our earlier conclusions, Fig. 9 shows that socially instructed learning is selected against if the learning schedule is shuffled or if individual learning precedes social learning. In these simulations, the proportion of social learning could not evolve to zero but instead, evolution tends to decrease the social learning rate to zero. That shows that, even if there is no trade-off between exploration and exploitation (number of learning episodes is fixed) and no trade-off between social and individual learning (proportion of social learning is fixed), socially-guided learning is detrimental and the time allocated to social learning is not used to adjust the network at all as social learning rate is zero. In replicates that evolve higher social learning rates in this condition, the performance of the networks is clearly lower.

**Fig. 9.**
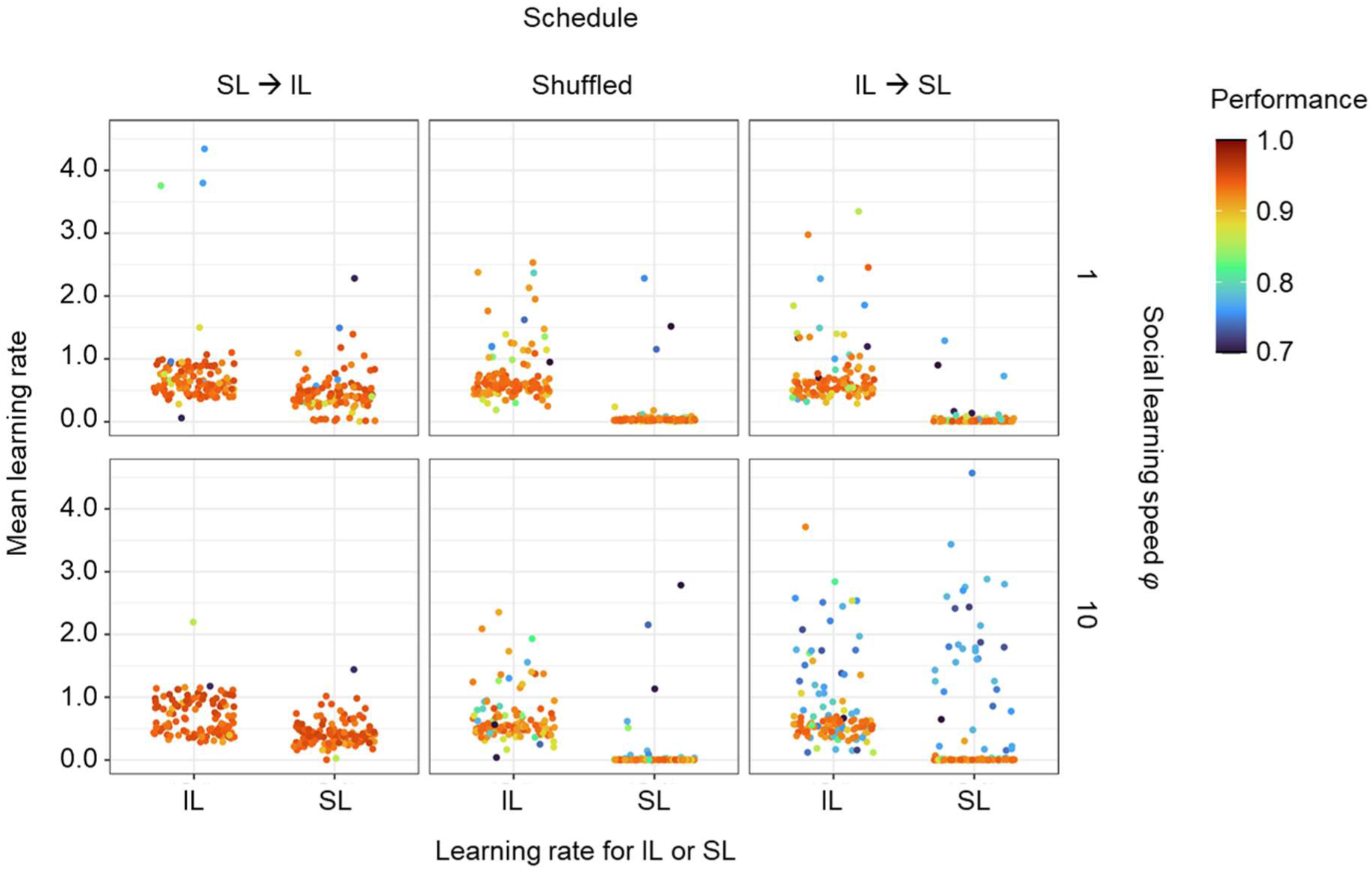
Comparison between social and individual learning rates in relation to the learning schedule and the speed of social learning. The plots show the rates of individual learning (IL) and social learning (SL) that evolved for different learning schedules (columns) and different social learning speeds (rows) in 60 replicate simulations. Each point represents the population mean averaged over the last 2K generations of the simulation. In all simulations, self-guided individual learning jointly evolved with socially instructed learning. Throughout the simulations, the number of learning episodes was fixed to 25 and the proportion of social learning was fixed to 0.25. Learning evolved under intermediate environmental stability (*f* = 0.1, *m* = 0.25, σ = 0.25).

For the SL→IL schedule, the rate of social learning is in most replicates higher than zero but lower than the rate of individual learning. This is to some extent surprising as in machine learning applications of neural networks it has been found to be favourable to start with a large learning rate and to reduce the learning rate as soon as high-fitness solutions have been found. However, in our model, social learning has a lower learning rate even though it is utilised first. This might be linked to the fact that social information is less reliable than the information gathered via individual learning and therefore the networks are adjusted to a lesser extent. However, this explanation is not in line with the surprisingly small effect of the speed of social learning: virtually the same rate of social learning evolved for a social learning speed of 1 as for a speed of 10 (although in the former case, social learning was lost in a larger number of cases). Since in this set of simulations the total number of time steps devoted to social learning is independent of its speed, one would expect a higher social learning rate when the speed of social learning is lower to compensate for the lower number of social learning events. Or when social learning speed is higher, the learning rate may be expected to be lower to take into account that social learning is not very reliable. Clearly, the effect of evolution on the learning rate needs further study.

## Discussion

We presented a novel modelling framework for studying the evolution of social learning in which the learning mechanism is explicitly taken into consideration. Our framework is based on a previous model by Kozielska and Weissing^20^, who used simple neural networks to investigate the evolution of individual learning in changing environments. We extended this model to consider social learning from more experienced demonstrators. The extended model uses the same evolvable neural network and the same learning mechanism (based on the Delta rule); social learning only differs from individual learning in the information used for learning. This way, different types of learning can be quite naturally addressed and combined in the same framework.

In this study, we focused on one form of individual learning and two forms of social learning: social guidance and social instruction. In both cases, a demonstrator points out instances that seem promising as test cases for learning, but the two forms differ in the feedback the learner receives when investigating such a test case. During social guidance, learners gather information by directly probing the environment; during social instruction, learners rely on the knowledge of their demonstrators. In the model, the neural network underlying learning, parameters like the learning speed and the learning duration, and the degree of reliance on social learning are heritable (subject to small mutations) and, therefore, evolve over the generations. This way, we could study whether and when subjects evolve to learn individually, socially, or via a combination of both forms of learning.

In line with^20^, we considered a learning task in which individuals had to infer the quality of items from cues in a world where the cue-quality relationship can change with environmental conditions. By systematically investigating different regimes of environmental change, we showed that the evolutionary outcome of our model agrees with intuitive expectations and previous literature (e.g.,^7,26,27,30–32^: any form of learning is selected against in very stable environments (where genetic adaptation is superior to learning), pure individual learning evolves in very unstable environments (where the knowledge of demonstrators gets readily outdated), and social learning (in combination with individual learning) evolves at intermediate levels of environmental stability. In our simulations, pure social learning could persist as a transient state for some time but was subsequently complemented or even replaced by individual learning after a while.

At intermediate levels of environmental stability, the evolved learning strategies depended strongly on the form of social learning. In the case of socially guided learning, populations readily evolved a high-performance combined learning strategy in which individual and social learning occurred in a random order. In contrast, socially instructed learning only evolved if the scheduling of social and individual learning could evolve as well. When this was the case, many simulations evolved towards a high-performance combined learning strategy where socially guided learning preceded individual learning. However, a substantial fraction of the replicate simulations evolved towards two alternative attractors corresponding to either pure individual learning or to the loss of learning.

Interestingly, occasional switches from one attractor to a different one occurred in many simulations involving socially instructed learning. Such switches between attractors are typical for stochastic systems with multiple attractors (see^33^ and references therein). Yet, to our knowledge, such switches have never been reported in the context of the evolution of learning strategies. Presumably, the more complex evolutionary dynamics observed in our study are related to the fact that our model has many more evolutionary degrees of freedom than most other models of the evolution of social learning. Many of our simulations did not converge to the nearest fitness peak in the most direct manner but instead explored a substantial part of the fitness landscape in the course of evolution.

Such ‘exploration’ could make it more likely that high-performance solutions are found when the model is applied to more complex and realistic scenarios^34^.

The differences in the evolution of socially guided and socially instructed learning highlight the importance of addressing the mechanism of learning when studying the evolution of social learning. Most previous models (e.g., those reviewed in ^26^) did not consider the evolution of a learning mechanism but directly described the effects of social and individual learning on the level of adaptation of the population. In these models, individual and social learning are typically represented by one or two parameters. For instance, individual learning is represented by the degree to which the true state of the environment is approximated or by the probability that an ‘adequate’ behaviour is chosen; social learning is represented by the fidelity with which learners copy the behaviour of their demonstrators. In other words, the learning process is not incorporated or addressed in these models.

In our model, the kind of information and the feedback available to the learners defines the different types of learning. In addition, since all types of learning rely on the same underlying mechanism, individuals can integrate the information they obtain via the different types of learning. During each learning step, individuals locally modify some of their neural connections based on the available information. The newly acquired information can then constitute the starting point for further learning.

In the case of socially guided learning, our results come the closest to the predictions of Boyd and Richerson^27,35^. At intermediate levels of environmental change, and with relatively lower costs of social learning, these authors showed that combining individual and social learning should always be favoured over pure individual learning. The analyses of Boyd and Richerson have often been interpreted as prescribing a learning schedule in which social learning precedes individual learning, even though this order did not matter much in their original model formulation^26^. Similarly, our results suggest that a social-learning-first schedule is not essential for evolving a highly adaptive combined learning strategy. Moreover, such a combined strategy evolved even though socially guided learning was as costly as individual learning. However, for other forms of social learning, the scheduling and costs of individual and social learning may be crucial for the evolutionary outcome. In our model, this was the case for socially instructed learning, where social learning could only get off the ground in combination with a social-learning-first schedule. It has been argued^7,30,36,37^ that a learning schedule in which social learning precedes individual learning may be adaptive because learners can first absorb the nongenetically encoded adaptive information acquired by previous generations and subsequently augment this information by individual experience. Models showed that such a learning schedule can indeed evolve and increase population performance^30^, but this only holds if specific requirements are met, such as a reduction of the learning time through a combination of social and individual learning^38,39^. In our model, evolved socially instructed learning in combination with and scheduled before individual learning did (marginally) increase average performance relative to pure individual learning, although the total number of learning episodes (and, hence, the costs of learning) was roughly the same (or even slightly higher) for the combination of social and individual learning as for pure individual learning.

Most classical approaches to social learning do not model the learning process explicitly (e.g. those reviewed in^26^). If they do, they often assume that ‘cultural traits’ are directly transmitted from a demonstrator to a learner. This has been criticised^10–12^ because the mental representations of ‘cultural traits’ are not replicated from a demonstrator to a learner, but rather are ‘reconstructed’ by the learner through an inferential process that is strongly affected by cognitive structures and processes. The social learning of language is a good example. Language exists in two distinct forms: external linguistic behaviours consisting of verbal utterances and internal linguistic representations stored in individual brains. When being taught a language, learners are exposed to the utterances of their demonstrators; from this, they have to distil the meaning and compositional rules of these utterances and incorporate them internally into their knowledge of the language. Therefore, many models of language acquisition and language evolution include this constructive process on the side of the learners (e.g.^40^). In contrast, few models of social learning and cultural evolution explicitly address this inferential process.

In our opinion, a neural network approach is ideally suited for representing the reconstructive process of knowledge acquisition via social learning. Even in the absence of a memory (as in our variant of a social-learning model), knowledge can be acquired by a learner – not by copying memory content but by changing the weights of the neural connections that embody the learner’s knowledge about the world. In other words, socially acquired information is not replicated but transformed and integrated into prior knowledge. This implies that the ‘learning outcome’ depends not only on the ‘learning content’ but also on the prior state of the learner’s network. Consequently, the order in which learning content is provided may strongly affect the final learning outcome. Similarly, the effectiveness of a demonstrator may depend on the ‘knowledge state’ of the learner, and different demonstrators may be most effective regarding knowledge transfer at different times in the learning process. In addition to the learning history, the learning outcome may also be affected by the structural features of the network. In our model, these include the network architecture and the genetically determined connection weights. It is conceivable that, similar to cognitive biases^15,41^, such structural features canalise the learning process and strongly determine the outcome^42^. In our model, these features are subject to genetic evolution. Accordingly, we would expect that they will, evolve to a state that makes the learning process more effective^20,43^.

Until now, there have been very few attempts to model the social transmission of information as a ‘reconstructive’ process. The recent modelling framework proposed by Falandays and Smaldino^44^ is perhaps the most promising one. The modularity of that framework is an attractive feature, as it allows to ‘plug in’ a variety of cognitive submodels. However, it includes some unrealistic ingredients (like Bayesian updating, cf.^41^) and it is less ‘tangible’ than the neural network approach proposed here. Making use of neural networks particularly allowed us to mechanistically account for how individuals acquire novel information and transform their knowledge in the course of social learning. While neural networks play an important role in machine learning, their use in understanding social learning and cultural evolution is surprisingly limited. This is even true in the field of evolutionary linguistics, which has long recognised that cultural transmission is a reconstructive process^23^ (but see^45^). Hutchins and Hazlehurst^46^ presented a pioneering model that provides proof of principle that a natural regularity (in their study: the relationship between moon phase and tidal state) that is too difficult to be unravelled by an individual can become known by a population via neural network-based social learning and cultural evolution. However, their model is specifically tailored to the question under study, does not include genetic evolution and learning is based on backpropagation, which is biologically unrealistic. Parisi, Nolfi, Acerbi and colleagues developed several neural network models to investigate social learning and cultural evolution in the context of foraging (e.g.^16,17,47^. These models have various ingredients in common with our approach (e.g., using the Delta rule for learning), but they include additional elements (e.g., the joint control of movement and foraging) that make it difficult to extrapolate their findings. Genetic evolution and coevolution between individual and social learning are not considered in these models. Gabora^18,48^ developed neural network models that are specifically tailored to the study of (the cultural evolution of) gestures and other movements used for communication or other tasks. These models are specifically tailored to the movement domain and do not account for knowledge transfer between generations or gene-culture coevolution. Borenstein and Ruppin^19^ and Curran and O’Riordan^49^ do allow for gene-culture coevolution, but their models focus on social learning, not considering the possibility of using individual learning. Moreover, these and other models (e.g.^50^), which are often inspired by machine learning, make use of sophisticated learning mechanisms that are removed from biological reality. Our approach incorporates many elements that, to our knowledge, have never been studied together: gene-culture coevolution, the joint evolution of individual and social learning, neural networks that can easily be adjusted to a variety of learning tasks, and a biologically inspired learning mechanism.

To allow comparison with previous work^20^, we focussed on a relatively simple learning task: locating the maximum of an environmentally determined function. This limits our conclusions in a variety of ways. Perhaps most importantly, all members of the population can achieve a similar ‘state of knowledge’ at the end of their learning period. As a consequence, the ability to select one of the best-performing individuals of the previous generation as a demonstrator (success bias) does not make much of a difference. Moreover, the learning task is not suited to study the evolution of cumulative culture. However, much more difficult learning tasks can easily be incorporated into our model framework. If the learning task is sufficiently sophisticated, such that very limited insight can be gained by individual learning (as the one considered by Hutchins and Hazlehurst^46^), social learning may unfold its full potential. In this case, the population members will, even at the end of the learning period, differ substantially in their knowledge, making demonstrator selection profitable. It is conceivable that different individuals gain expertise in different domains of knowledge, which may make knowledge transfer beneficial. If knowledge is spread in the population, one can imagine the emergence of subpopulations specialising in specific knowledge domains and the ‘personalised’ choice of a demonstrator from one or several of these subpopulations. In brief, the model might provide new perspectives on the evolution of cumulative culture.

On purpose, we considered a very simple feedforward network, but this is not a fundamental aspect of the model. The network structure can easily be expanded or adapted to other purposes. One could also allow the network architecture to evolve (by including ‘macro-mutations’ leading to the emergence of new network nodes or connections, or to the removal of existing nodes and connections). Perhaps most interestingly, one could let the ‘learning connections’ evolve, allowing the learning process to make optimal use of the network structure (see also Discussion in^20^).

In the field of cultural evolution, there is much debate on how cultural evolution, and the social transmission of information as its fundamental ingredient, should be modelled^9^. Members of the so-called ‘Paris School’ criticised current modelling approaches as they do not do justice to the fact that ‘information reconstruction’ is an important aspect of social learning (e.g.^10–12^). While members of the ‘California School’ agree that social learning is a reconstructive process, they have argued that the details of the knowledge acquisition process do not matter too much for the final outcome of social learning and cultural evolution (e.g.^13^). Whether this is indeed the case can only be judged once a broad spectrum of research questions is not only addressed by the standard approaches but also by ‘reconstructive’ learning frameworks like the one developed here.

## Methods

### Model setup

By means of individual-based simulations, we study the evolution of a population that lives in an environment with characteristics that are constant within generations but may change from one generation to the next. The basic setup of the model is based on our previous work^20^ on unguided learning. Briefly, individuals have a fixed lifetime divided into discrete timesteps. At the start of their life, they can spend a number of timesteps learning. During the learning period, individuals can use individual learning, social learning, or a mixture of both to adapt their inherited neural network to their current environment. After the learning period, individuals switch to foraging, using their modified (via learning) network to assess the available foraging options. During the foraging period, a number of food items are offered per timestep, of which an individual chooses the one that, according to their neural network, has the highest quality. The expected lifetime reproductive success of an individual is proportional to the sum of the qualities of all items chosen throughout the foraging period. In other words, this sum is a proxy for fitness in our model. Offspring inherit their parents’ neural networks and their learning strategy (the duration of the learning period, the learning rates, and the reliance on individual versus social information), subject to rare mutations.

Over the generations, the neural networks and the learning strategies tend to improve and adapt to the structure of the environment (the relationship between cues and quality and the rate and degree of change of this relationship).

### Environment

In our model, the environment contains a continuum of food items differing in quality *Q*, where *Q* ranges from 0 to 1. Food quality cannot be perceived directly, but only on the basis of cues like colour or smell. The relationship between cues *C* and quality *Q* is described by a Gaussian function (the environmental profile; see Supp. Fig. 1) with mode *μ* and standard deviation *σ*:

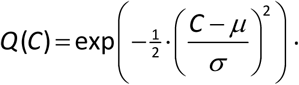

As a default, we used *σ* = 0.25, but other values are also investigated. Throughout the simulations, we restricted the set of cues to the interval [-1, 1] and we assumed that they could be arranged on a circle where -1 is connected to +1. This way, the environmental profile function was wrapped on this interval to keep the total amount of resources in the environment constant (see Supplementary Fig. 1).

The environmental parameters *μ* and *σ* are constant throughout a generation, but the mode *μ* of the environmental profile shifts randomly to the left or to the right at regular intervals (starting at *μ* = 0 in generation 0). The frequency *f* of environmental change is constant throughout each simulation; here, we report on the values *f* = 0.01, 0.1, and 1 (i.e., the mode shifts every 100, 10, or 1 generation, respectively). The magnitude of the mode shift was drawn from a normal distribution with mean *m* and standard deviation 0.05. *m* is constant throughout each simulation. Here, we report on the values *m* = 0.1, 0.25, and 0.4.

### Neural networks

Each individual possesses a neural network that is used to predict the quality of food items on the basis of their properties (cues). We consider relatively simple neural networks consisting of 10 neuron-like nodes (see Supplementary Fig. 2). Kozielska and Weissing^20^ showed that this network is able to learn and make efficient foraging decisions. A network *N* receives a cue *C* as input and produces an output *Q_N_* (*C*) that can be interpreted as the predicted quality of a food item emitting that cue. The networks consist of nodes that are organised in a sequence of layers (one input node, two hidden layers, each with four nodes, and one output node; see Supplementary Fig. 2). Each node (except the output) is connected to one or several nodes in the subsequent layer, and it can stimulate or inhibit the activities of these nodes. Each connection has a certain strength that is indicated by a ‘weighing factor’ *w*, where a positive value of *w* represents stimulation, while a negative value corresponds to inhibition. The input node receives the cue value *C* which is a real number. Hidden nodes and the output node have a baseline activation, *b,* and are further activated or inhibited by the nodes from the previous layer. The activity of the hidden nodes is constrained between 0 and 1 by the clamped ReLU function and the output of the network can be any real number (for more details see^20^ and Supplementary Fig. 2).

All weighing factors and baseline activations of nodes are heritable and transmitted from parents to offspring (subject to mutation, see below) and are therefore subject to evolution. Additionally, some weighing factors can change during the individual’s lifetime via learning.

### Learning: the Delta rule

Learning is conducted in discrete, usually multiple episodes at the beginning of an agent’s lifespan. In the default version of the model, one learning event occurs during each learning episode. In each learning event, the learner inspects one food item emitting a certain cue, say *C_I_*. If *L* is the learner’s neural network, the learner will predict that the item has quality *P* = *Q_L_* (*C_I_*). Subsequently, the learner receives feedback on the correctness of this prediction by comparing it with a ‘target value’ *T*. This target value depends on the type of learning considered (see Fig. 1). For example, *T* can be the ‘true’ quality of the food item: *T* = *Q* (*C_I_*). The discrepancy Δ = *T* − *P* between target value and predicted quality is then used to modify the final four weighing factors of the neural network (see Fig. S2) by making use of the ‘Delta rule’:

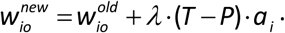

Here, 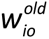 is the weighing factor of the connection between node *i* in the preceding layer and output node *o* before the learning event. The new value 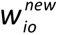 is obtained by adding the term *λ* (*T* – *P*)·*a_i_* 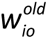, where *T* − *P* is the discrepancy between target value and prediction, *a*_i_ is the activation level of node *i*, and *λ* is the ‘learning rate’, which is a heritable factor that determines how strongly the weighing factors are modified during each learning event. We allow the learning rate to differ between individual and social learning (see below).

As argued in^20^, this form of neural network-based learning has three important advantages: (1) it is biologically realistic; (2) it is efficient (at least for the learning task considered here); and (3) it is simple in that learning is ‘localised’ at a few connections (rather than restructuring the whole neural network as in backpropagation and other machine-learning algorithms).

### Four types of learning

#### Unguided individual learning

Per learning event, the learning item is selected at random. The corresponding random cue *C_R_* induces the learner to predict the quality as *P* = *Q_L_* (*C_R_*). The feedback received (the target value) is the true quality of the item: *T* = *Q* (*C_R_*). This type of learning is implemented in^20^ and corresponds to an individual exploring the environment in a trial-and-error fashion and directly learning the value of cues. Unguided learning lacks focus in the sense that all kinds of cues (and not only the promising ones) are used for learning. Depending on the characteristics of the environment, this can have advantages and disadvantages.

#### Self-guided individual learning

Per learning event, the ‘most promising’ item on offer is chosen as the learning item. To be more precise, ten random items are presented to the learner, and the learner chooses the cue 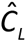 that it predicts to be associated with the highest quality. The predicted quality of the chosen item is therefore: 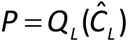. The target value is again the true value of the chosen item: 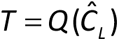. This type of learning focuses on the presumed peak of the environmental profile, thus neglecting less promising parts of the profile. This may make learning more efficient, but it still bears the risk that the presumed maximiser 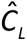 may not be close to the true maximiser of the environmental profile *Q*(*C*).

#### Socially guided learning

Per learning event, the demonstrator points out the most promising item on offer. Again, ten random items are presented to the learner, and the demonstrator points out the cue 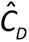 that the demonstrator’s network predicts to be associated with the highest quality. To the learner, the predicted quality of this item is 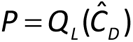. The target value is again the true value of the chosen item: 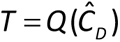. In this form of social learning, which is often called ‘stimulus enhancement’ [REFs], the demonstrator has the sole role of guiding the learning process.

#### Socially instructed learning

The cue chosen for learning is again the cue 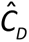 considered to be most promising by the demonstrator. The value predicted by the learner is again 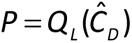. However, now the target value is not the true quality of the item 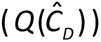 but the item quality as predicted by the demonstrator’s network: 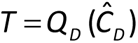. Hence, this type of learning is equivalent to teaching and the ‘true’ quality of the chosen item 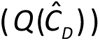 is not checked by the learner, which implies that the learning outcome can be suboptimal if the demonstrator’s predicting profile *Q_D_* (*C*) does not match the environmental profile *Q*(*C*) very well. However, the lack of a reality check has the advantage that socially instructed learning can proceed faster and with fewer costs than the other types of learning.

All four types of learning and their effect on the learner’s profile (the expected relationship between cues and quality) are illustrated in Figure 1. Simulations always allowed just one of the individual learning implementations and one of the social learning implementations to coevolve.

As illustrated in Fig. 1, we consider four distinct types of learning. These differ from each other in two aspects: the way the item to be learnt from is chosen and the way the target value is determined. The first two types of learning are variants of individual learning: they only depend on the learner and the learner’s neural network *L*. The other two types of learning are (at least partly) based on social information and are therefore variants of social learning. In these types of learning, a ‘demonstrator’ with neural network *D* plays an important role. Throughout, we assume that the demonstrator is an individual of the previous generation who has undergone learning in that generation. We considered various ways to select a demonstrator, which will be explained in a later section.

#### Socially instructed learning

The cue chosen for learning is again the cue 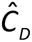 considered to be most promising by the demonstrator. The value predicted by the learner is again 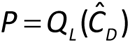. However, now the target value is not the true quality of the item 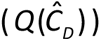 but the item quality as predicted by the demonstrator’s network: 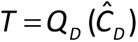. Hence, this type of learning is equivalent to teaching and the ‘true’ quality of the chosen item 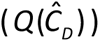 is not checked by the learner, which implies that the learning outcome can be suboptimal if the demonstrator’s predicting profile *Q_D_* (*C*) does not match the environmental profile *Q*(*C*) very well. However, the lack of a reality check has the advantage that socially instructed learning can proceed faster and with fewer costs than the other types of learning.

All four types of learning and their effect on the learner’s profile (the expected relationship between cues and quality) are illustrated in Fig. 1. Simulations always allowed just one of the individual learning implementations and one of the social learning implementations to coevolve.

### Default settings and extensions

#### Choice of demonstrators

By default, the demonstrators for social learning are chosen at random from the individuals of the previous generation (after having completed their learning and foraging period). For each learning event, a new demonstrator is chosen. Social learning may be more effective if particular individuals are chosen as demonstrators^28,29^. Therefore, we also implemented a variant of social learning where the choice of demonstrator is based on the demonstrator’s ‘success’, which we call a ‘success bias’. In this variant, demonstrators are no longer picked completely at random by learners. Instead, the probability of becoming a demonstrator is proportional to an individual’s fitness (the sum of qualities of the food items collected by the individual during its foraging period).

#### Speed of learning

By default, our model assumes that there is one learning event per timestep. As explained above, socially instructed learning could be ‘cheaper’ than the other forms of learning because it is based on the demonstrator’s knowledge and does not require a reality check. To account for this, we considered a speed advantage of socially instructed learning by including multiple learning events from the same demonstrator within one timestep. The number of learning events per timestep is indicated by the speed parameter *φ*. By default, *φ* = 1, but we also considered *φ* = 10.

#### Learning strategies

The ‘learning strategy’ of an individual specifies the duration *δ* of the learning period, the type(s) of learning employed during the learning period (individual vs. social learning), the rates of individual and social learning (*λ_i_* and *λ_s_*, see below) and in some cases the learning schedule (see below). In all our simulations, the individuals have a fixed lifespan of 500 time units. The duration of the learning period is an integer *δ* (satisfying 0 ≤ *δ* < 500) that corresponds to the number of timesteps spent learning at the start of an individual’s life. The remaining 500 − *δ* timesteps are spent foraging.

In each timestep spent learning only one type of learning can be employed. However, we allow the evolution of combined forms of learning, where one type of individual learning (in this study, this is always self-guided learning) can be combined with one type of social learning. It is determined at the start of each simulation which type of social learning (socially guided or socially instructed) is combined with self-guided learning. Individual and social learning can either be combined in a random order (‘shuffled learning’) or in a predetermined order (‘scheduled learning’). We allow for the evolution of different learning rates for both types of learning: a value *λ_i_* used when employing individual learning and a value *λ_s_* used when employing social learning.

#### Shuffled learning

At the start of each timestep spent learning, the learning type employed in that timestep will be drawn at random, where *π_s_* is the probability that social learning will be employed while *π_i_* = 1 − *π_s_* is the probability that individual learning will be chosen. Hence, the expected number of timesteps spent on individual learning is *π_i_* ·*δ*, while *π_s_* ·*δ* is the expected number of timesteps spent on social learning. *π_s_* (0 ≤ *π_s_* ≤ 1) is a heritable parameter. If *π_s_* evolves to very small values, evolution leads to ‘pure individual learning’, while ‘pure social learning’ evolves if *π_s_* evolves to a value close to 1 (see Fig. 3A). Unless specified otherwise, shuffled learning was used in the simulations.

#### Scheduled learning

Now, either individual learning precedes social learning (‘IL→SL’) or social learning precedes individual learning (‘SL→IL’). The type of schedule is either predetermined at the start of a simulation or is the outcome of evolution (see below). In the context of scheduled learning, the parameter *π_i_* has a slightly different interpretation than in shuffled learning: *π_i_* ·*δ*, rounded to the next integer value, is the number of timesteps spent on individual learning, while *π_s_* ·*δ*, rounded to the next integer value, is the number of timesteps spent on social learning.

#### Evolution of shuffle or schedule

Our final heritable parameter χ takes on a discrete value to represent the learning schedule: this corresponds to either shuffled learning, the schedule SL→IL, or the schedule IL→SL. Via rare mutations (see below), χ can evolve to any of the three possible states.

### Reproduction and inheritance

#### Reproduction

Following learning and foraging for all individuals, synchronous reproduction occurs. The expected reproductive success of an individual is proportional to the individual’s ‘fitness’, the sum of the qualities of the food items collected by the individual during its foraging period. For simplicity, we assume that the population size *N* is constant and that reproduction is asexual. For each of the *N* individuals of the offspring generation, a parent is drawn with replacement and the probability that a given individual is drawn as a parent is proportional to the individual’s fitness. Following reproduction, the individuals of the parental generation survive to act as demonstrators for social learning in the offspring population. Afterwards, all individuals in the parental generation die.

#### Inheritance

Our model includes the following heritable parameters: the 24 weighing factors associated with the connections of the neural network and the 9 baseline activations of the nodes of the network (excluding the input node); the duration *δ* of the learning period; the learning rates *λ_i_* and *λ_s_* in the cases of individual and social learning; the (expected) proportion *π_s_* of timesteps spent on social learning; and the parameter *χ* determining the learning schedule. For simplicity, we assume that the individuals of our model are haploid and that each heritable parameter is represented as an allele at a specified gene locus. All these alleles except for *χ* correspond to numerical values. Hence, the allelic value 15 at the *δ* locus instructs the individual to spend 15 timesteps on learning (and 485 timesteps on foraging). During reproduction, the parental allele at any locus can mutate, with a mutation probability of 0.01 per locus. When a mutation occurs at a gene locus representing a real-valued model parameter (all parameters except *δ* and *χ*), the parental allele *A* is changed to *A* + *ε*, where the mutational step size *ε* is sampled from a normal distribution with mean zero and standard deviation 0.1. When a mutation occurs at the *δ* locus (which represents an integer), the mutational step size is *ε* = ±1, both options occurring with equal probability. Finally, when a mutation occurs at the *χ* locus (which represents the three options: shuffled learning, SL→IL, or IL→SL), any of the two alternative options is drawn with equal probability.

By the interplay of selection, mutation, and genetic drift, all heritable parameters of the model can evolve. However, specific scenarios can be investigated by switching off mutation at specific loci. For example, shuffled learning and the two forms of scheduled learning can be investigated in isolation of each other by assigning the corresponding state of *χ* to all members of the initial population and by setting the mutation probability at the *χ* locus to zero.

### Simulation setup

All simulations reported were run with a population size of *N* = 1000 individuals. All evolvable parameters (i.e., all parameters that were not kept fixed by setting the mutation rate to zero) were initialised by assigning an allelic value to each individual of the initial population. In the case of the weighing factors and baseline activation levels of the neural network, the initial allelic values were drawn from a uniform distribution on the interval [-1,+1]; initial learning rates were drawn from a uniform distribution on the interval [0,1], the initial duration of the learning period was set to *δ* = 20, the initial proportion of social learning was set to *π_s_* = 0.5, and *χ* was initialised to a shuffled schedule.

Most simulations were run for 20 thousand generations, but evolutionary equilibrium (as judged by the achieved performance) was usually reached in a much shorter time. The simulation code was written in C++ and run on the Peregrine and Habrok High-Performance Computing clusters of the University of Groningen, the Netherlands.

### Data availability

Examples of simulation output files and summary data used for the figure are available at https://dataverse.nl/privateurl.xhtml?token=bb201313-0b86-41f4-9510-6c8d780d1b57.

### Code availability

The source code (in C++) for the simulation program is available at https://dataverse.nl/privateurl.xhtml?token=bb201313-0b86-41f4-9510-6c8d780d1b57.

## Acknowledgements

We thank the MARM group members and especially Stefano Tiso for valuable discussions. This work was supported by the European Research Council (ERC Advanced Grant No. 789240 awarded to F.J.W.). J.C. was supported by the Mobility European Master in Evolution (MEME) programme.

## Author contributions

All authors contributed to the design of the study. M.K. and J.C. wrote the simulation code. J.C. analysed the results. All authors were closely involved in discussing of the results and the preparation of the manuscript.

## Additional information

Supplementary Figs 1-12 can be found in the Supplementary file.

**Correspondence** and requests for materials should be addressed to Magdalena Kozielska or Franz J. Weissing.

