## Supplementary Figures 1-12 for "A neural network model for the evolution of reconstructive social learning"

**Jacob Chisausky, Ines Daras, Franz J. Weissing, and Magdalena Kozielska**

This file includes the following supplementary figures:

Supplementary Figure 1. Illustration of the relationship between cues and environmental quality.

Supplementary Figure 2. A schematic of the neural network used in this study.

Supplementary Figure 3: Evolution of unguided and self-guided individual learning in relation to the characteristics of the environment.

Supplementary Figure 4: Lifetime energy gain ('fitness') achieved by evolved unguided and self-guided individual learning.

Supplementary Figure 5: Joint evolution of self-guided individual learning and socially guided learning in relation to the characteristics of the environment.

Supplementary Figure 6: Joint evolution of unguided individual learning and socially guided learning in relation to the characteristics of the environment.

Supplementary Figure 7: Joint evolution of self-guided individual learning and socially instructed learning in relation to the characteristics of the environment.

Supplementary Figure 8: Joint evolution of unguided individual learning and socially instructed learning in relation to the characteristics of the environment.

Supplementary Figure 9: Joint evolution of self-guided individual learning and socially instructed learning with success bias, higher social learning speed and an evolving learning schedule.

Supplementary Figure 10: Joint evolution of unguided individual learning and socially instructed learning with success bias, higher social learning speed and an evolving learning schedule.

Supplementary Figure 11: Fitness achieved in different regions of the fitness landscape in Figure 8.

Supplementary Figure 12: Long-term joint evolution of self-guided individual learning and socially instructed learning.

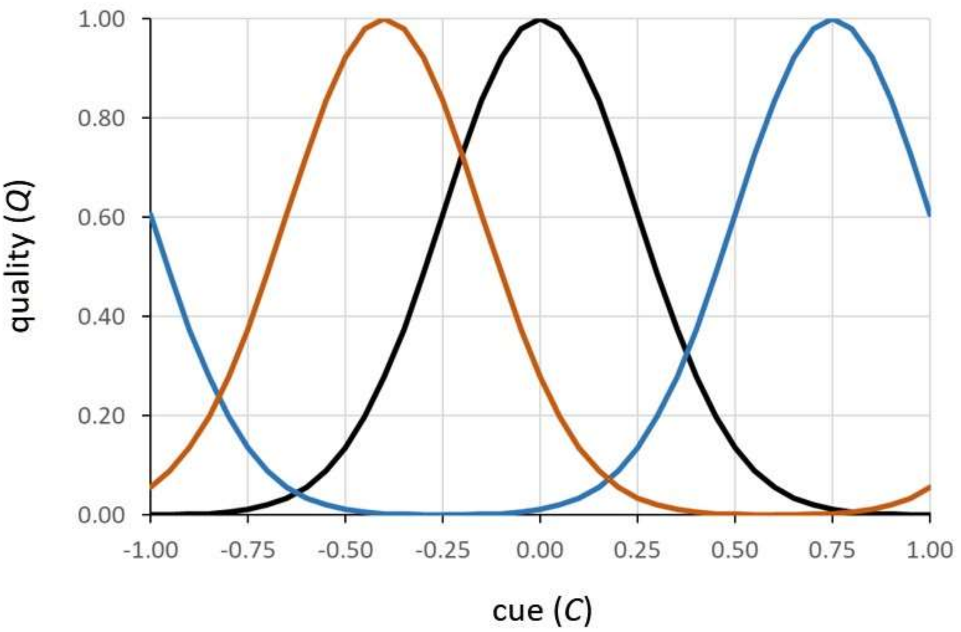


**Supplementary Figure 1. Illustration of the relationship between cues and environmental quality.** Three Gaussian functions illustrate the relationship between cues ($C$) and quality ($Q$) in three different generations. The peak of the quality function can shift between generations, indicating a change in the environment. Each colour shows the environment at a different time. Here, $\sigma = 0.25$.

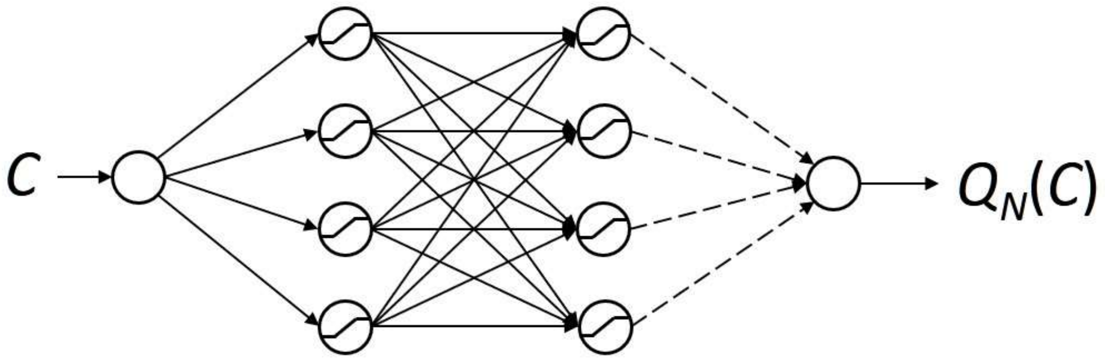


**Supplementary Figure 2. A schematic of the neural network used in this study.** Our network receives a cue *C* as input, and it produces an output $Q_N(C)$, which can be interpreted as the predicted quality of an item emitting that cue. In this model, we use a network with one input and one output (C and $Q_N(C)$, respectively) and two hidden layers, each with four nodes. Arrows indicate the information flow in the network. Each node (except for the output node) is connected to one or several nodes in the subsequent layer, and it can stimulate or inhibit the activities of these nodes. Each connection has a certain strength - weight *w*, where a positive value of *w* represents stimulation, while a negative value corresponds to inhibition. The input node receives the cue value *C* which is a real number. This value is processed and determines the node activities at the subsequent level. More precisely, the activity $y_i$ of node *i* in each layer is given by an expression of the form $y_i = A\left(\sum_j w_{ij} x_j + b_i\right)$. Here *j* runs over all nodes of the previous layer that are connected to *i*, $x_j$ is the activity of node *j*, and $w_{ij}$ is the strength of the connection between nodes *j* and *i*. $b_i$ is the baseline activation of node *i*. Function *A* is a so-called "activation function." In this paper, we used the "clamped ReLU" function that is fast and returns 0 for arguments lower than 0, and 1 for arguments larger than 1. For arguments between 0 and 1, it returns the argument without transformation. No activation function was used for the output node. Solid arrows represent genetically hardwired connections that do not change during learning. Dashed arrows represent the genetically determined weights that can also change during learning (see the main text for details).

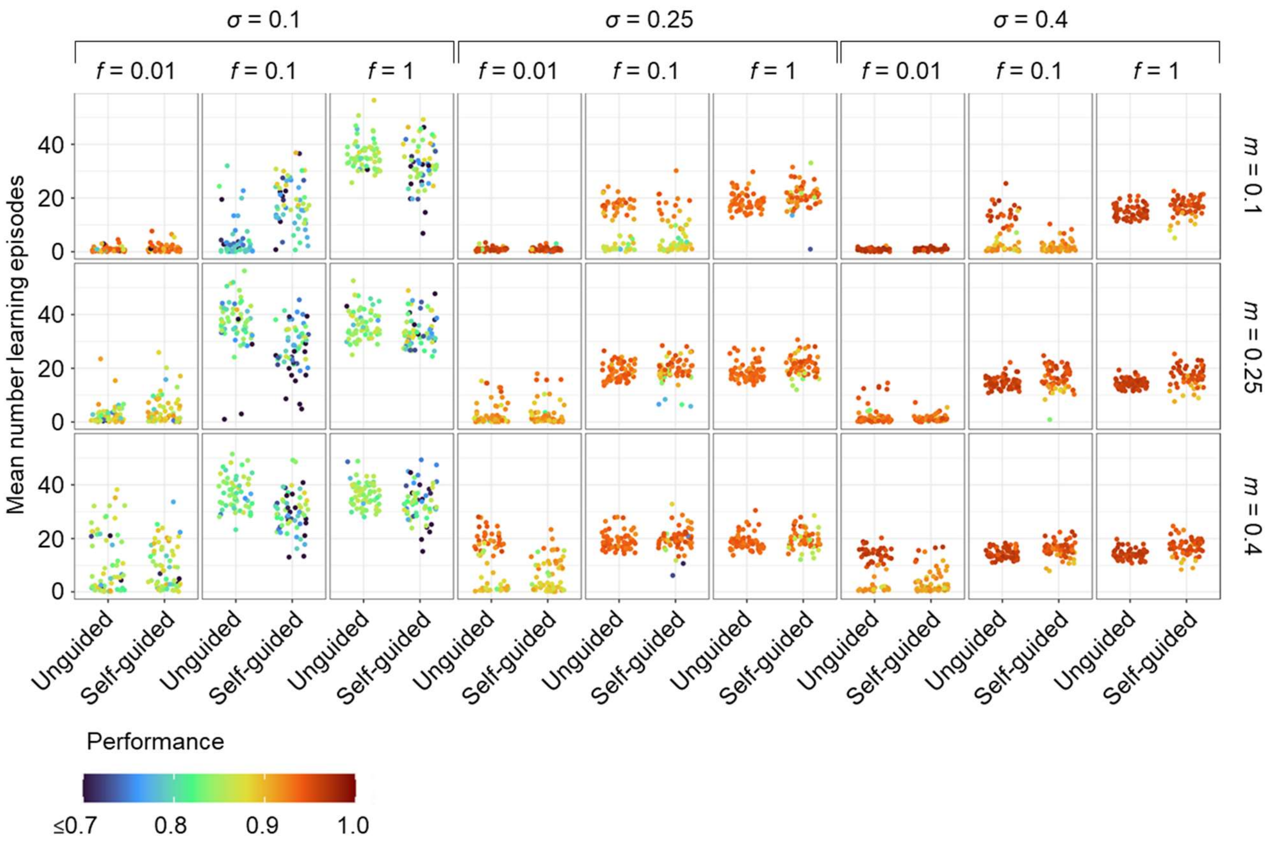


**Supplementary Figure 3**: **Evolution of unguided and self-guided individual learning in relation to the characteristics of the environment.** This figure expands Figure 2 in the main text by considering various values of $\sigma$, the width of the environmental profile. The simulation setup and the plotting conventions are as in Figure 2. This figure shows that the width of the environmental profile has a relatively small effect on whether learning evolves, however when it does the number of learning episodes tends to increase with decreasing $\sigma$. At large $\sigma$, unguided and self-guided learning evolve in the same environmental conditions, with only a few exceptions: self-guided learning evolves (the number of learning episodes is greater than 0 in most replicates) for the scenario with $\sigma$=0.1, $m$=0.1 and $f$=0.1. In that case, exploring the environment randomly does not seem to support the evolution of learning in some environments that lead to the evolution of self-guided learning The opposite is true for $\sigma$=0.4, $m$=0.1 and $f$=0.1 where in most replicates self-guided learning does not evolve. This can stem from the fact that the performance achieved by this type of learning is relatively lower and may not compensate for the time spent learning in this environment.

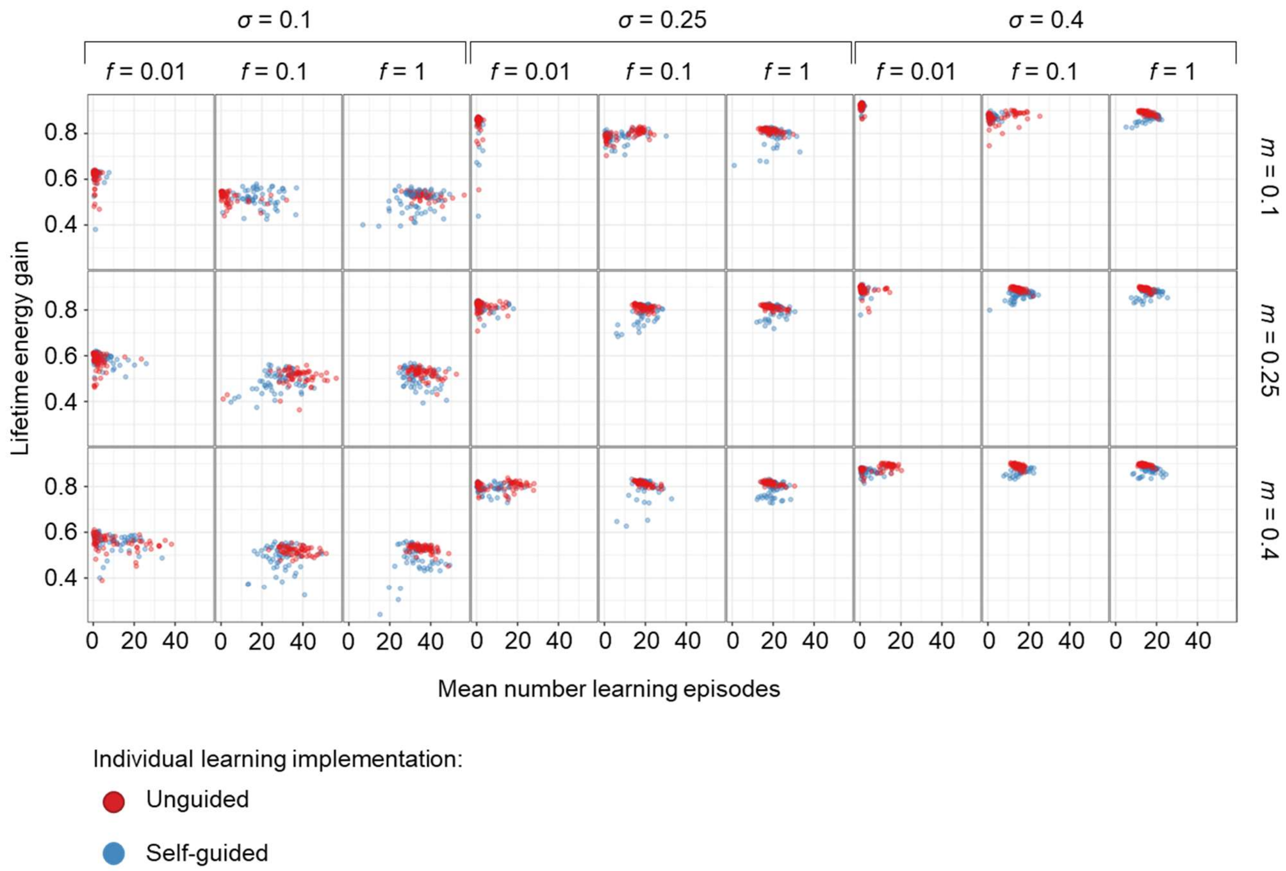


**Supplementary Figure 4. Lifetime energy gain ('fitness') achieved by evolved unguided and self-guided individual learning.** For the simulations in Supp. Fig. 3, the lifetime energy gain (our measure of fitness) is plotted against the evolved number of learning episodes. Each point corresponds to a replicate, with a mean number of learning episodes and lifetime energy gain averaged over all individuals and the last 2K generations of evolution. The colour of the points indicates simulations with unguided (red) and self-guided (blue) individual learning. From the figure, we conclude that populations using unguided learning tend to have somewhat higher fitness than the ones using self-guided learning. However, in both cases, average fitness increases with increasing $\sigma$ as the amount of resources in the environment increases.

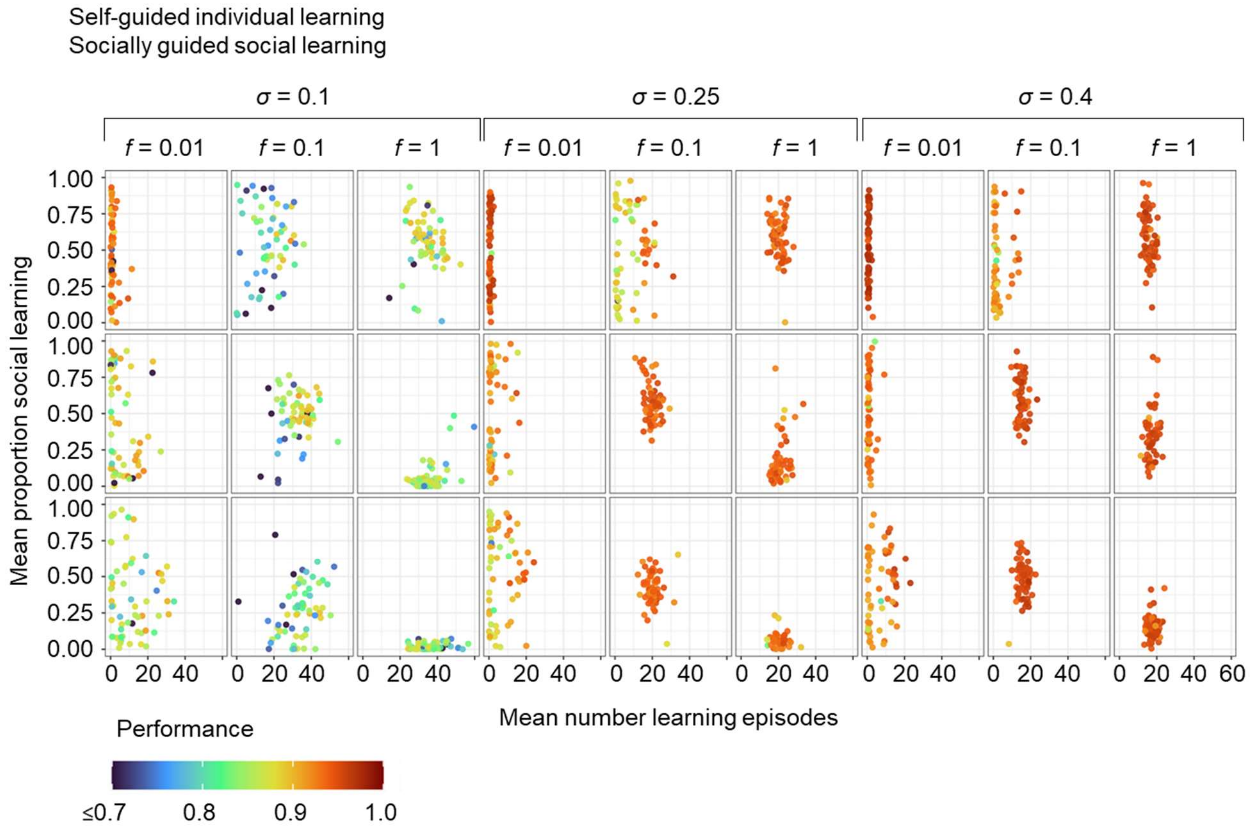


**Supplementary Figure 5**: **Joint evolution of self-guided individual learning and socially guided learning in relation to the characteristics of the environment**. This figure expands Figure 4 in the main text by considering various values of *σ*, the width of the environmental profile. The simulation setup and the plotting conventions are as in Figure 4. From the expanded figure we conclude that the width of the environmental profile (*σ)* has a relatively weak effect on the evolution of social learning with chances of social learning evolving increasing with increasing *σ* for the fast environmental change (*f*=1). The effects of other environmental variables (m and f) are discussed in the main text.

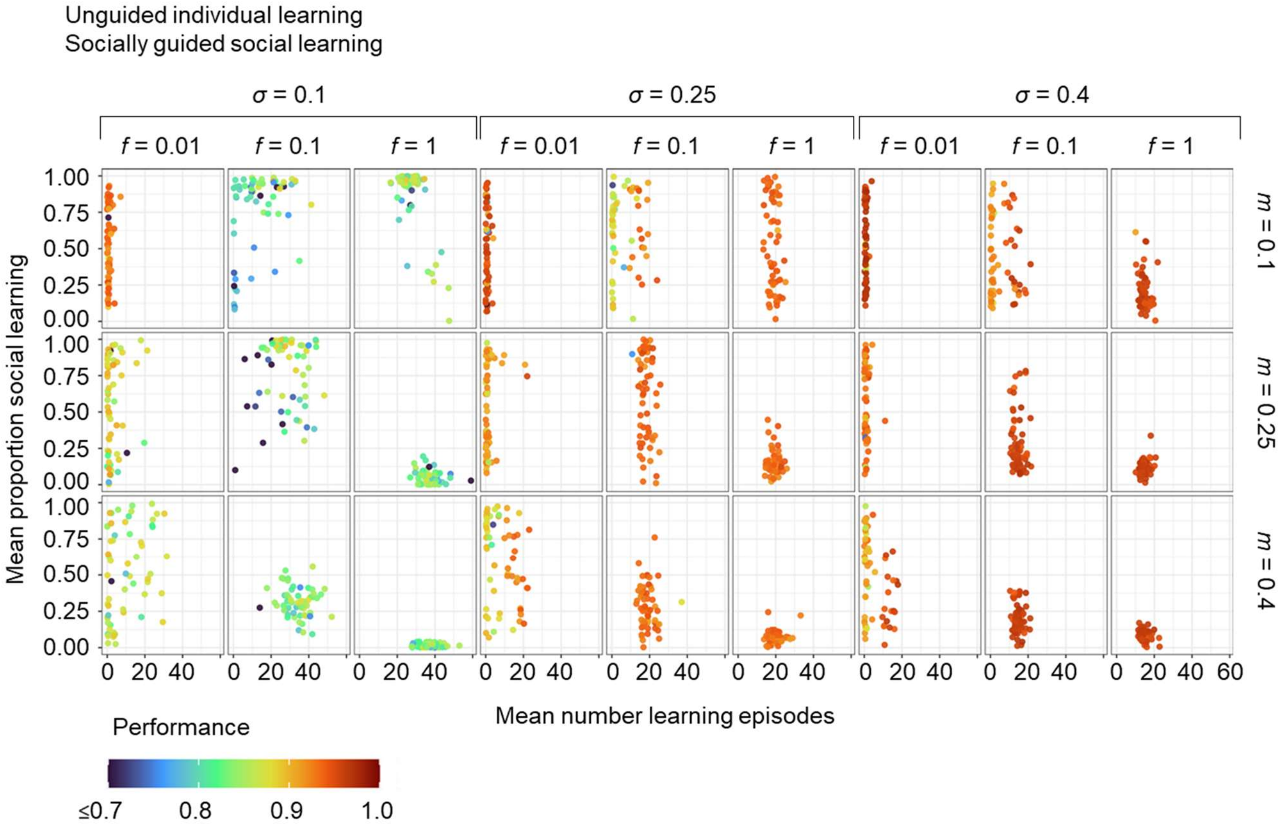


**Supplementary Figure 6**: **Joint evolution of unguided individual learning and socially guided learning in relation to the characteristics of the environment**. This figure is a counterpart to Supp. Fig. 5: the simulation setup and the plotting conventions are as in Supp. Fig. 5, but self-guided individual learning is replaced by unguided individual learning. The comparison of Supp. Figs 5 and 6 reveals that social learning evolves in the same environmental conditions, independent of the type of individual learning it coevolved with. However, when social learning evolves, the proportion of social learning is more variable between the replicate simulations when evolving with unguided individual learning. In that case, some populations rely exclusively on social learning (mean proportion of social learning is near 1).

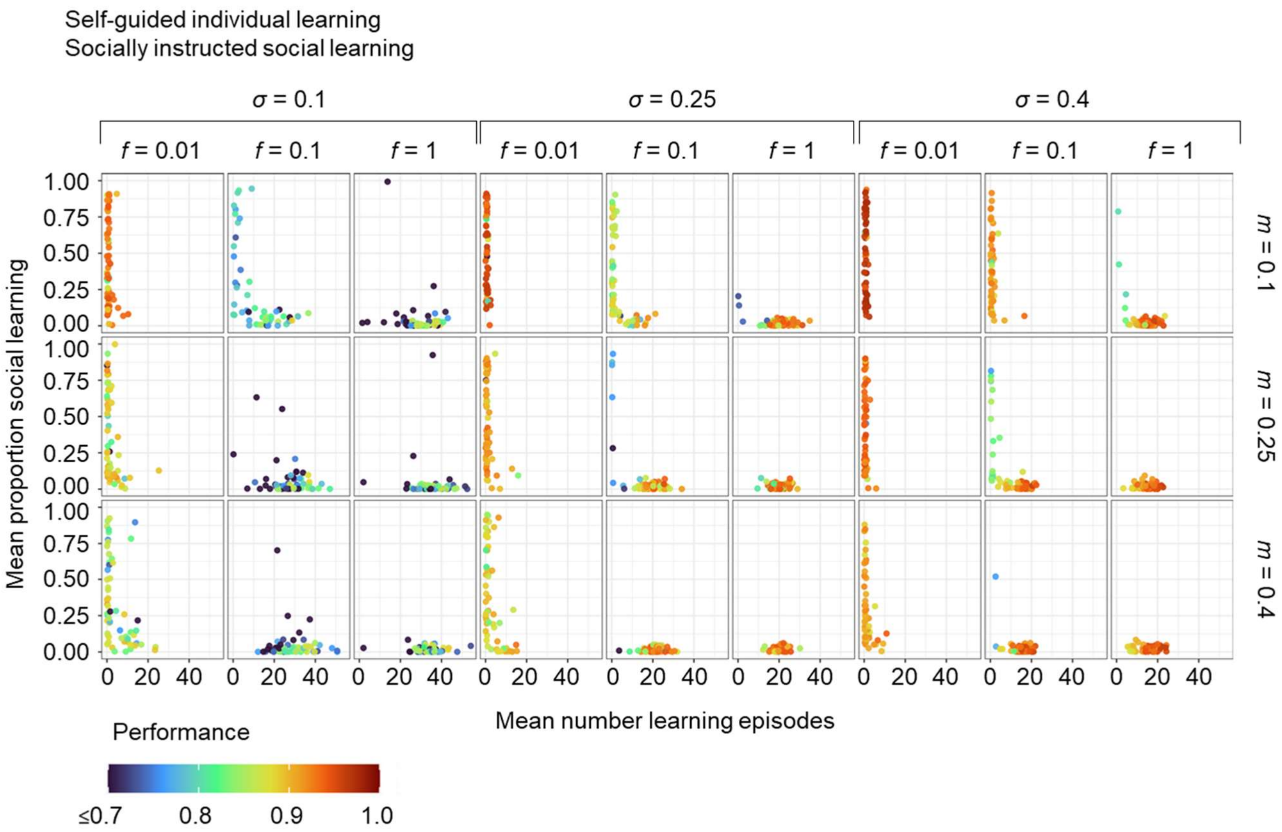


**Supplementary Figure 7**: **Joint evolution of self-guided individual learning and socially instructed learning in relation to the characteristics of the environment**. This figure expands Figure 5 in the main text by considering various values of $\sigma$, the width of the environmental profile. The simulation setup and the plotting conventions are as in Figure 5. From the expanded figure we conclude that socially instructed social learning does not evolve in any of the environments considered. Some exceptional replicates with a considerable proportion of social learning and a positive number of learning episodes have low fitness indicating that the presence of social learning is not a good strategy.

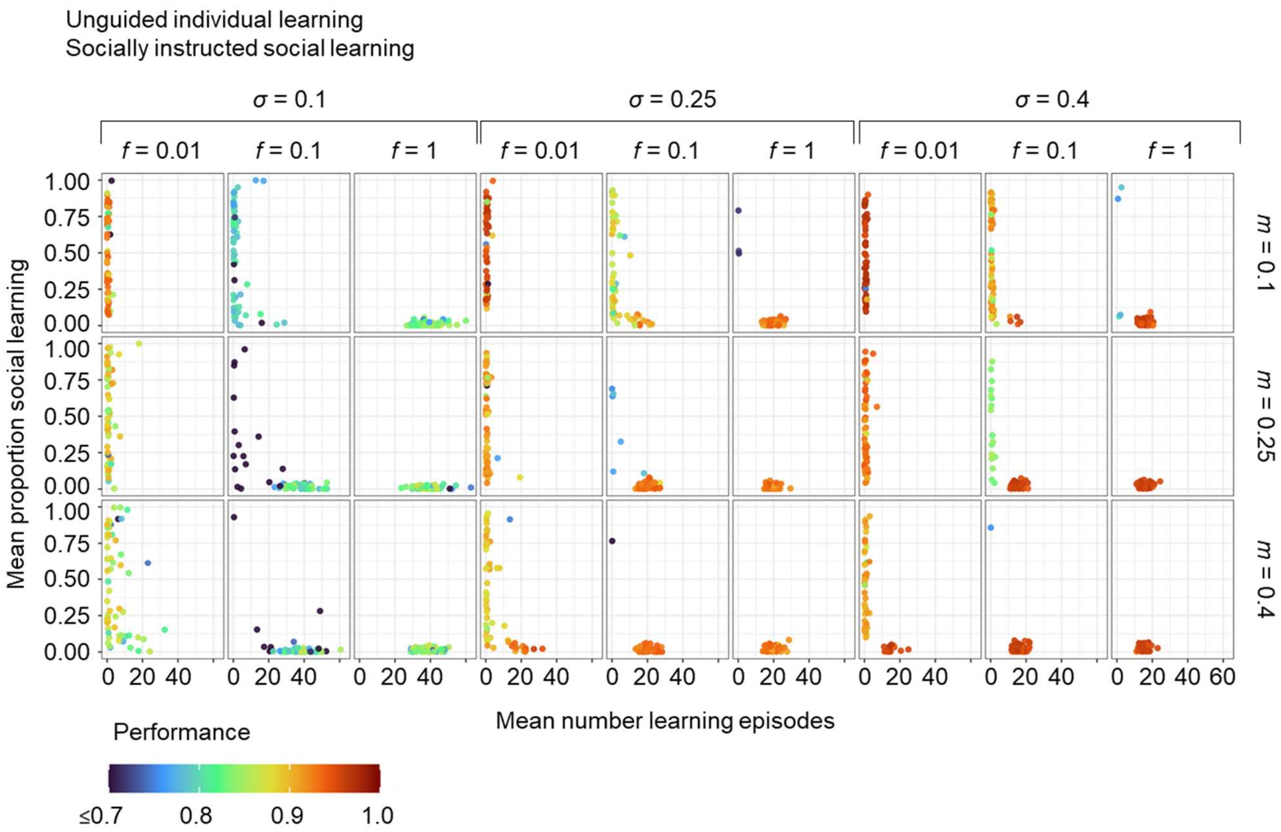


**Supplementary Figure 8**: **Joint evolution of unguided individual learning and socially instructed learning.** This figure is a counterpart to Supp. Fig. 7: the simulation setup and the plotting conventions are as in Supp. Fig. 7, but self-guided individual learning is replaced by unguided individual learning. The similar results of Supp. Figs 7 and 8 reveals that socially instructed learning does not evolve, independent of the type of individual social learning present.

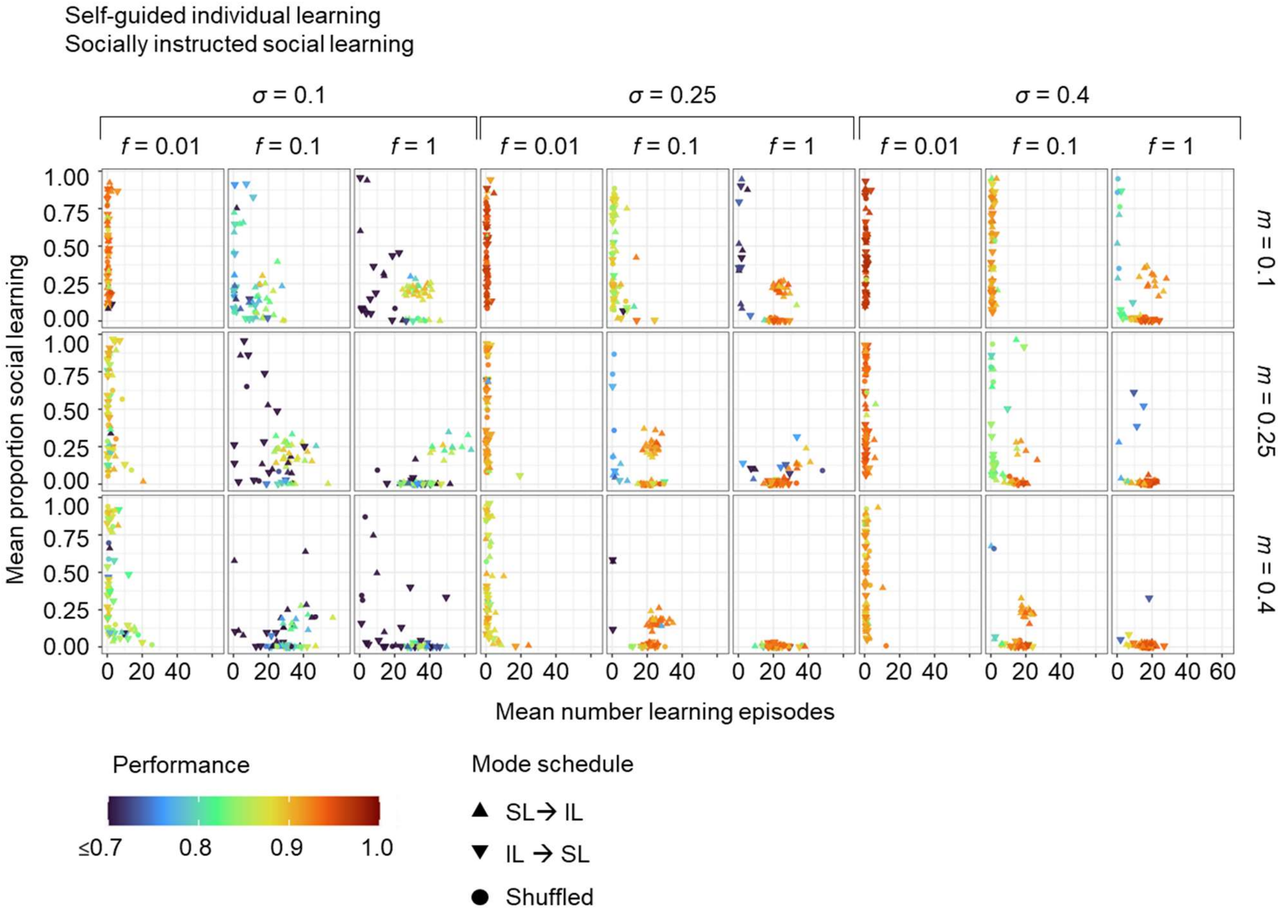


**Supplementary Figure 9**: **Joint evolution of self-guided individual learning and socially instructed learning with success bias, higher social learning speed and an evolving learning schedule**. This figure expands Figure 7 in the main text by considering various values of $\sigma$, the width of the environmental profile. The simulation setup and the plotting conventions are as in Figure 7. From the expanded figure we conclude that socially instructed learning can evolve also for narrower and wider environmental profiles. However, social learning is less likely to evolve for the wide environmental profile ($\sigma$=0.4). For the narrow environmental profile ($\sigma$=0.1) many populations evolve social learning, but only the ones with SL→IL schedule can achieve relatively high performance.

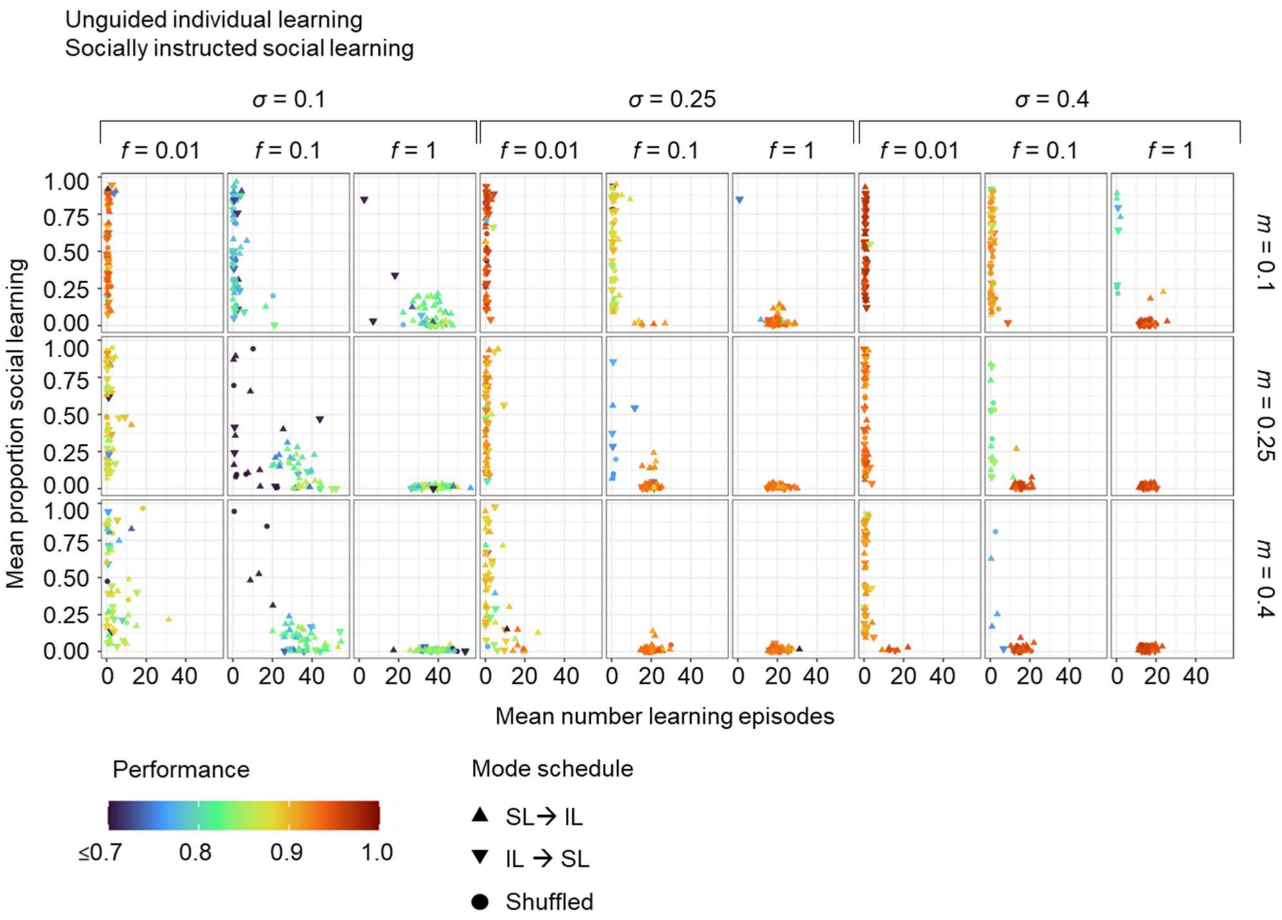


**Supplementary Figure 10**: **Joint evolution of unguided individual learning and socially instructed learning with success bias, higher social learning speed and an evolving learning schedule**. This figure is a counterpart to Supp. Fig. 9: the simulation setup and the plotting conventions are as in Supp. Fig. 9, but self-guided individual learning is replaced by unguided individual learning. The comparison of Supp. Figs 9 and 10 reveals that socially instructed learning is less likely to evolve when unguided social learning is utilised, except for some conditions with narrow environmental profile.

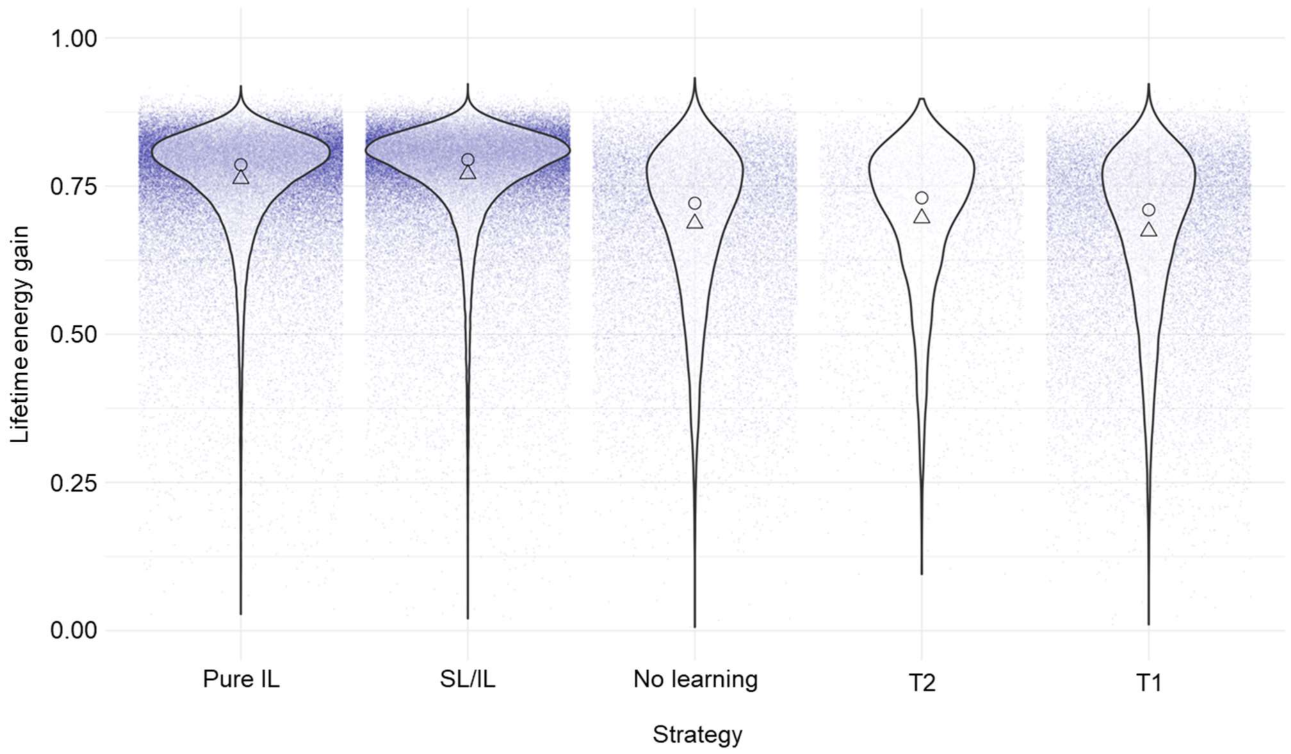


**Supplementary Figure 11: Fitness achieved in different regions of the fitness landscape in Figure 8 of the main text.** For 1000 replicate simulations, the average fitness of the evolving population is plotted every 101$^{th}$ generation (after an initial period of 2000 generations) in relation to the region in phase space that is visited by the evolutionary trajectory at that time. The five regions are indicated in Fig. 8 of the main text. Noise is added to the x-axis to increase readability. Violins are scaled so that each violin has the same area. Triangles show means of the data and circles show medians.

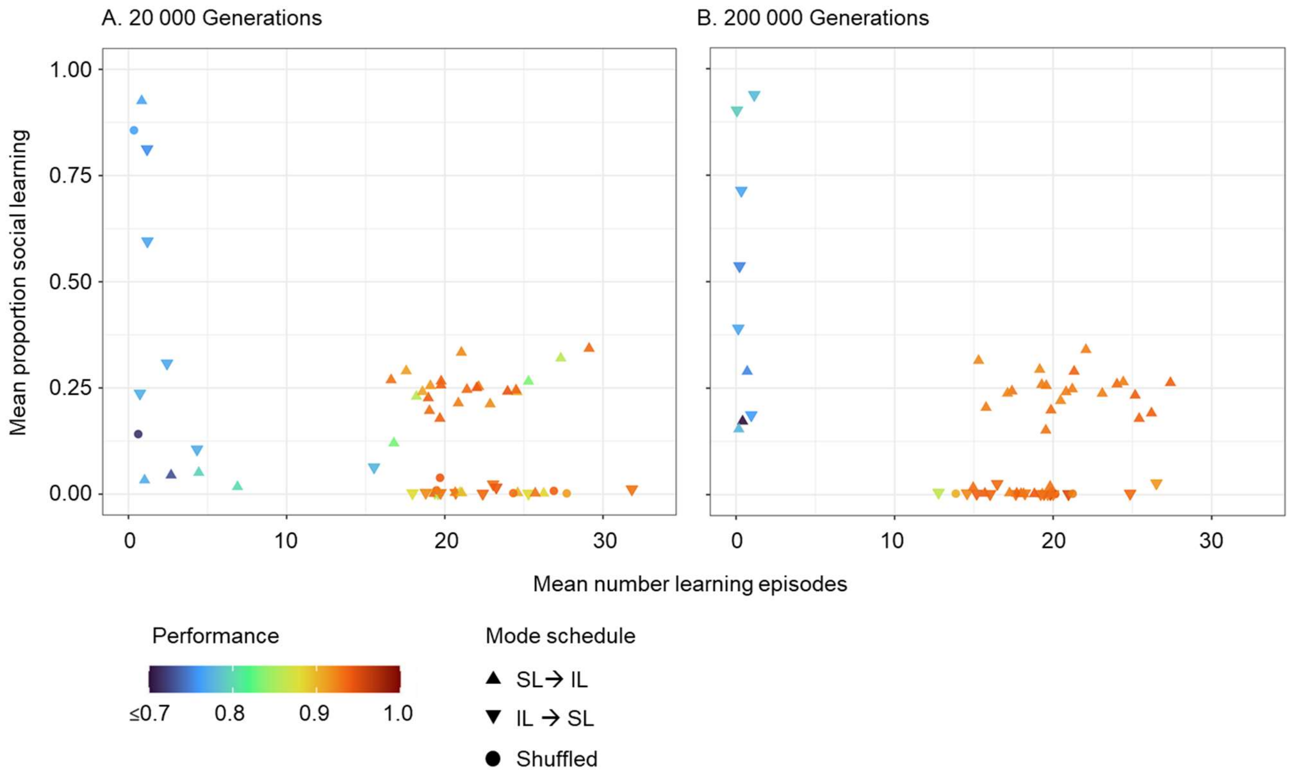


**Supplementary Figure 12: Long-term joint evolution of self-guided individual learning and socially instructed learning.** Under the same conditions as in Fig. 8 (and the middle panel of Figure 7 in the main text); population mean values of 60 replicate simulations are displayed after **(A)** 20K generations and **(B)** 200K generations. Even after a 10x longer simulation time than our default value of 20K, the same three 'attractors' are still present in the population: no learning, pure individual learning, and a combination of individual and social learning.
